# Enhanced FGFR3 activity in post-mitotic principal neurons during brain development results in cortical dysplasia and axon miswiring

**DOI:** 10.1101/2020.05.02.073924

**Authors:** Jui-Yen Huang, Bruna Baumgarten Krebs, Marisha Lynn Miskus, May Lin Russell, Eamonn Patrick Duffy, Jason Michael Graf, Hui-Chen Lu

**Author notes:** Corresponding Author and Address: Hui-Chen Lu, Ph.D., Department of Psychological and Brain Sciences, Indiana University, 1101 E. 10th Street, Bloomington, IN 47405, Jui-Yen Huang, Ph.D., Department of Psychological and Brain Sciences, Indiana University, 1101 E. 10th Street, Bloomington, IN 47405.

## Abstract

Abnormal levels of fibroblast growth factors (FGFs) and FGF receptors (FGFRs) have been detected in various neurological disorders. The potent impact of FGF-FGFR in multiple embryonic developmental processes makes it challenging to elucidate their roles in post-mitotic neurons. Taking an alternative approach, we directly examined the impact of aberrant FGFR function after neurogenesis by generating a FGFR gain-of-function (GOF) transgenic mouse which expresses constitutively activated FGFR3 (FGFR3^K650E^) in post-mitotic glutamatergic neurons. We found that enhanced FGFR activity in glutamatergic neurons results in abnormal radial migration and axonal miswiring. Regarding the lamination phenotype in GOF brains, we found later-born Cux1-positive neurons are dispersed throughout the GOF cortex. Such a cortical migration deficit is likely caused, at least in part, by a significant reduction of the radial processes normally projecting from the radial glia cells (RGCs). In addition, FGFR3 GOF also results in the misrouting of several long-range axonal projections, including the corpus callosum, anterior commissure, and postcommissural fornix. RNA-sequencing analysis of the GOF embryonic cortex reveals significant alterations in several pathways involved in cell cycle regulation and axonal pathfinding. Collectively, our results suggest that FGFR hyperfunction in post-mitotic neurons at the late embryonic stage result in cortical dysplasia and circuit miswiring.

## Introduction

The fibroblast growth factors (FGFs)-FGF receptors (FGFRs) family is comprised of structurally related tyrosine kinases receptors (FGFR1-4) and 22 FGF ligands ^1,2^. These receptors interact with a large group of FGF ligands and initiate several different intracellular signaling pathways to regulate diverse biological processes ^2,3^. Several FGFs/FGFRs are abundantly expressed in neuroprogenitors during early brain development ^2,4–7^. Compelling studies demonstrate that FGFs-FGFRs are key regulators of neurogenesis, neural differentiation, and cortical patterning in embryonic development ^2,7–11^. During early mouse embryonic development, neocortical neurons are generated from embryonic day 10.5 (E10.5) when neuroepithelial progenitors begin to divide symmetrically to expand their numbers and differentiate into the radial glial cell (RGCs) that initiate neurogenesis ^12,13^. It has been demonstrated that *Fgf10* promotes the transition of neuroepithelial cells to become RGCs ^11^. Additionally, RGCs also act as stem cells that divide in an increasingly asymmetric manner to self-renew and generate restricted intermediate progenitor cells (IPCs) and neurons ^14^. In this step, FGFR signaling in RGCs inhibits the transition from RGCs to IPCs and maintains the self-renewal ability of RGCs ^4,15^. Taking into consideration that the important function of FGF-FGFR signaling in governing neurogenesis, diminishing FGF-FGFR signaling results in loss of cortical surface area and a reduced number of glutamatergic neurons ^5,15–18^. Conversely, a constitutively active FGFR3 mutation results in increased neuronal number and larger brain size ^19,20^. In the late stage of cortical development (around E17.5 in mice), RGCs switch from the neurogenic phase to a gliogenic phase ^21–23^, a recent study further demonstrates that FGF-FGFR signaling is necessary and sufficient to redirect cell fate from neurons to astrocytes ^24^.

During early postnatal brain development, neuronal FGFRs are required to mediate neural activity-dependent processes in shaping dendritic patterning of cortical glutamatergic neurons ^25,26^. FGF7 and FGF22 differentially regulate inhibitory and excitatory synapse formation in the hippocampus ^27,28^. Taken together, FGFRs in post-mitotic neurons are required for dendritic patterning and synaptogenesis. However, the role of FGF-FGFR signaling in cell-type specific differentiation of new-born neurons has yet to be explored. Here we employed the NEX-Cre line ^29^ to express *FGFR3*^*K650E*^ (a constitutively active FGFR3) mainly in post-mitotic glutamatergic neurons to bypass the impact of FGF-FGFR in neurogenesis and to elucidate FGFR3’s roles in neuronal differentiation and neural circuit wiring. Results from this gain-of-function (GOF) approach reveal that the strong potent impacts of FGFR3 hyperfunction on cortical and hippocampal lamination, brain size, neuronal differentiation, and axonal pathfinding.

## Results

### Expressing FGFR3^K650E^ in NEX-lineage post-mitotic neurons results in aberrant cortical lamination

To explore the role of FGF-FGFR signaling in cortical neurons after neurogenesis, we expressed a constitutively activated FGFR3 mutant, *FGFR3*^*K650E*^, in post-mitotic glutamatergic neurons. Specifically, we generated a FGFR gain-of-function (GOF) mouse model by crossing the transgenic mouse line carrying a *FGFR3*^*K650E*^ conditional allele, CAG-flox-stop-flox-*FGFR3*^*K650E*^-IRES-eGFP ^30^ with the NEX-Cre mouse line ^29^. The progenies carrying both NEX-Cre and *FGFR3*^*K650E*^ conditional alleles (abbreviated as GOF mice) express FGFR3^K650E^ in cortical and hippocampal glutamatergic neurons (Fig. 1A). FGFR activation can result in the phosphorylation on FGFR substrate α (FRS2α) and extracellular signal-regulated kinase 1/2 (ERK1/2) ^2^. Western blotting of cortices prepared from postnatal day 7^th^ (P7) control and GOF mice reveals a significant increase in the abundance of phosphorylated FRS2α (p=0.0402) and ERK1/2 (p=0.0015), normalized to total FRS2α and ERK1/2, in GOF cortex compared to their littermate controls verifying an increase of FGFR3 signaling (Fig. 1B, C). These data verified an increase of FGFR3 signaling in the GOF cortex.

**Figure 1.**
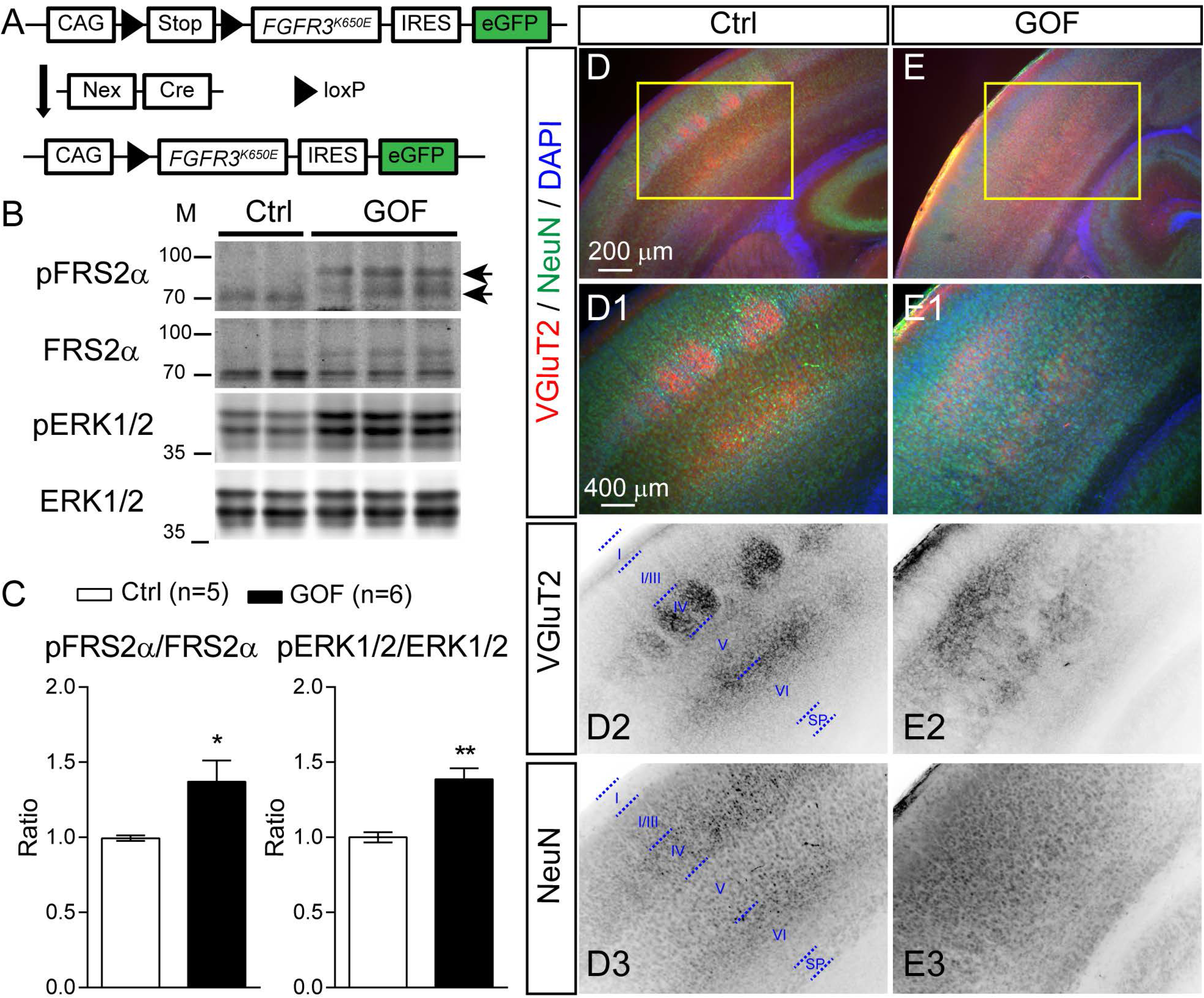
Expressing FGFR3^K650E^ in NEX-lineage neurons results in aberrant cortical lamination. (A) The diagram shows how GOF mice were generated. (B) Western blots show the abundance of phosphorylated FGFR substrate 2 α (FRS2α-Tyr196; pFRS2α), FRS2α, phosphorylated ERK1/2 (Thr202/Tyr204; pERK1/2), ERK1/2 in the S1 cortex of P7 control (Ctrl) and GOF mice. (C) Summaries for the fold changes in pFRS2α to FRS2α and pERK1/2 to ERK1/2 (Ctrl, n=5; GOF, n=6) in GOF mice. (D-E) VGluT2, NeuN and DAPI staining in the P7 S1 cortex. D1-3 and E1-3 show the enlarged images in D and E (yellow boxes). D2-3 and E2-3 show the inverted images of individual channels. I-VI, cortical layers; SP, subplate. Student’s-t test. *, p < 0.05; ***, p < 0.001. See Supplementary Figure 7 for full-length blots.

The mouse primary somatosensory (S1) cortex is known for its distinctive whisker-related patterns, called barrels. Barrel formation requires coordinated development between the innervating thalamocortical axons and the migrating layer IV neurons. Barrel pattern can be disrupted by defective neuronal differentiation/migration, axonal pathfinding ^23,31,32^. To visualizing the whisker-related patterns, we conducted immunostaining by using antibodies against NeuN (neuronal marker) and vesicular glutamate transporter 2 (VGluT2, thalamocortical axons marker) ^33–35^ in P7 GOF and littermate controls. In control brains, VGluT2 signals are evident as whisker-related patches in cortical layer IV and a distinctive band in layer VI (Fig. 1D, D1, D2). However, the distribution of VGluT2 signals in the GOF S1 cortex was dispersed throughout the middle of the cortical plate and lacked a distinctive pattern (Fig. 1E, E1, E2). Additionally, different from the lamination pattern revealed by NeuN in the control S1 cortex (Fig. 1D3), the distribution of NeuN in the GOF S1 cortex is rather uniform (Fig. 1E3).

Different from the typical ear-drop shape seen in control hippocampi (Supplementary Fig. 1A), GOF hippocampi are more rounded in shape (Supplementary Fig. 1B). Instead of the compacted pyramidal layers seen in control hippocampi (Supplementary Fig. 1C, C1), NeuN^+^ cells in GOF hippocampi were rather dispersed (Supplementary Fig. 1D, D1). Additionally, an ectopic cluster of neurons was often found directly above the CA1 region of GOF hippocampi (Supplementary Fig. 1B, D, D1). The neurons in the hilus of the dentate gyrus area were also dispersed more widely in GOF hippocampi (Supplementary Fig. 1F, F1), compared with control (Supplementary Fig. 1E, E1). The thickness of both the CA1 stratum pyramidale (p=0.0014) and the hilus (p<0.0001) were significantly greater in GOF compared to control mice (Supplementary Fig. 1G, H). Taken together, FGFR3 hyperfunction in glutamatergic neurons perturbs the lamination in the cortex and hippocampus. The presence of VGluT2 verifies that thalamocortical axons reached S1 cortex of GOF mice

### Postnatal FGFR3 GOF did not perturb cortical laminations

To determine the time window when FGFR3 GOF impact cortical lamination, we generated postnatal GOF mice by crossing the mice carrying CAG-flox-stop-flox-*FGFR3*^*K650E*^-IRES-eGFP allele with Nex-Cre(ERT2) mice and activated Cre by injecting tamoxifen after birth (Supplementary Fig. 2A; abbreviated as postnatal GOF mice). Specifically, these time-specific GOF pups and littermate controls were subjected to 100 mg/kg tamoxifen via single intraperitoneal injection at P1. The significant increases in the abundance of phosphorylated FRS2α (p=0.0136) and ERK1/2 (p=0.0373), normalized to total FRS2α and ERK1/2 respectively, in the P7 cortex from postnatal GOF pups (Supplementary Fig. 2B, C) confirmed the Cre-mediated recombination and upregulation of *FGFR3*^*K650E*^ in postnatal GOF pups. Immunofluorescent staining showed that GFP was expressed in the cortex and hippocampus of P7 postnatal GOF mice (Supplementary Fig. 2E, E1), but not control animals (Supplementary Fig. 2D, D1), again confirming appropriate Cre activation.

Cux1 is a homeodomain-containing DNA binding protein that is abundantly expressed in later-born cortical glutamatergic neurons ^36,37^. In both control (Supplementary Fig. 2D, D2) and postnatal GOF mice (Supplementary Fig. 2E, E2), Cux1^+^ neurons are distributed throughout layer II-IV and are enriched in layer IV barrels. DAPI staining also revealed distinctive lamination and barrels in both control and postnatal GOF mice (Supplementary Fig. 2D, D3, E, E3). Taking together the embryonic GOF data, we find that FGFR3 hyperfunction in post-mitotic glutamatergic neurons perturbs cortical lamination during late embryogenesis (between E12 and birth) but not after birth.

### Expressing FGFR3^K650E^ in NEX-lineage principal neurons resulted in misplacement of later-born, but not earlier born principal neurons

Lamination abnormalities are often resulted from radial migration deficits ^38–40^. To further explore if the lamination abnormality is due to misplacement of specific neuronal subtypes, we performed immunostaining with antibodies against cortical layer-specific markers Cux1, special AT-Rich Sequence-Binding protein 2 (Satb2), and COUP-TF-interacting protein 2 (ctip2) ^36,37,41^. In P7 GOF cortex, later-born Cux1^+^ cells were dispersed throughout the cortical plate (Fig. 2B, B1), quite different from their upper layer localization found in control (Fig. 2A, A1). Such misplacement of Cux1^+^ neurons in the GOF cortex was already evident by E18.5 (Fig. 2C, C1, D, D1). In GOF cortex, there were significantly fewer Cux1^+^ cells in the cortical plate (bin 8, p<0.0001; bin 9, p<0.0001), while more were present in the intermediate zone (IZ) and subventricular zones (SVZ) (bin 4, p=0.0415; bin 5, p=0.0057; bin 6, p<0.0001)(Fig. 2E). There was no difference in the total number of Cux1^+^ cells between control and GOF mice (p=0.6122; Fig. 2F). To further examine the distributions of different types of cortical neurons, Satb2 and Ctip2 double immunostaining was conducted (Fig. 2G, G1-2, H, H1-2). We found significantly more Satb2^+^ cells in GOF brain distributed in deep layers (bin 2, p<0.0001; bin 3, p<0.0001), towards the ventricular zone (VZ) (Fig. 2H, H1). Ctip2^+^ marks early-born glutamatergic neurons and they are often found in the deep layer ^36^. Distinct from Cux1^+^ or Satb2^+^ cells, the distribution of Ctip2^+^ cells is normal in the GOF cortex (Fig. 2G2, H2). The number of Satb2^+^ (p=0.2685; Fig. 2K) and Ctip2^+^ (p=0.3486; Fig. 2L) cells is similar between GOF and control mice. However, the percentage of Satb2 and Ctip2 double-positive cells is significantly higher in the GOF cortex (p=0.005; Fig. 2M). Taken together, these results suggest that FGFR3 GOF in principal neurons disrupts the migration of layer II-IV cortical neurons and alters neuronal subtype specification.

**Figure 2.**
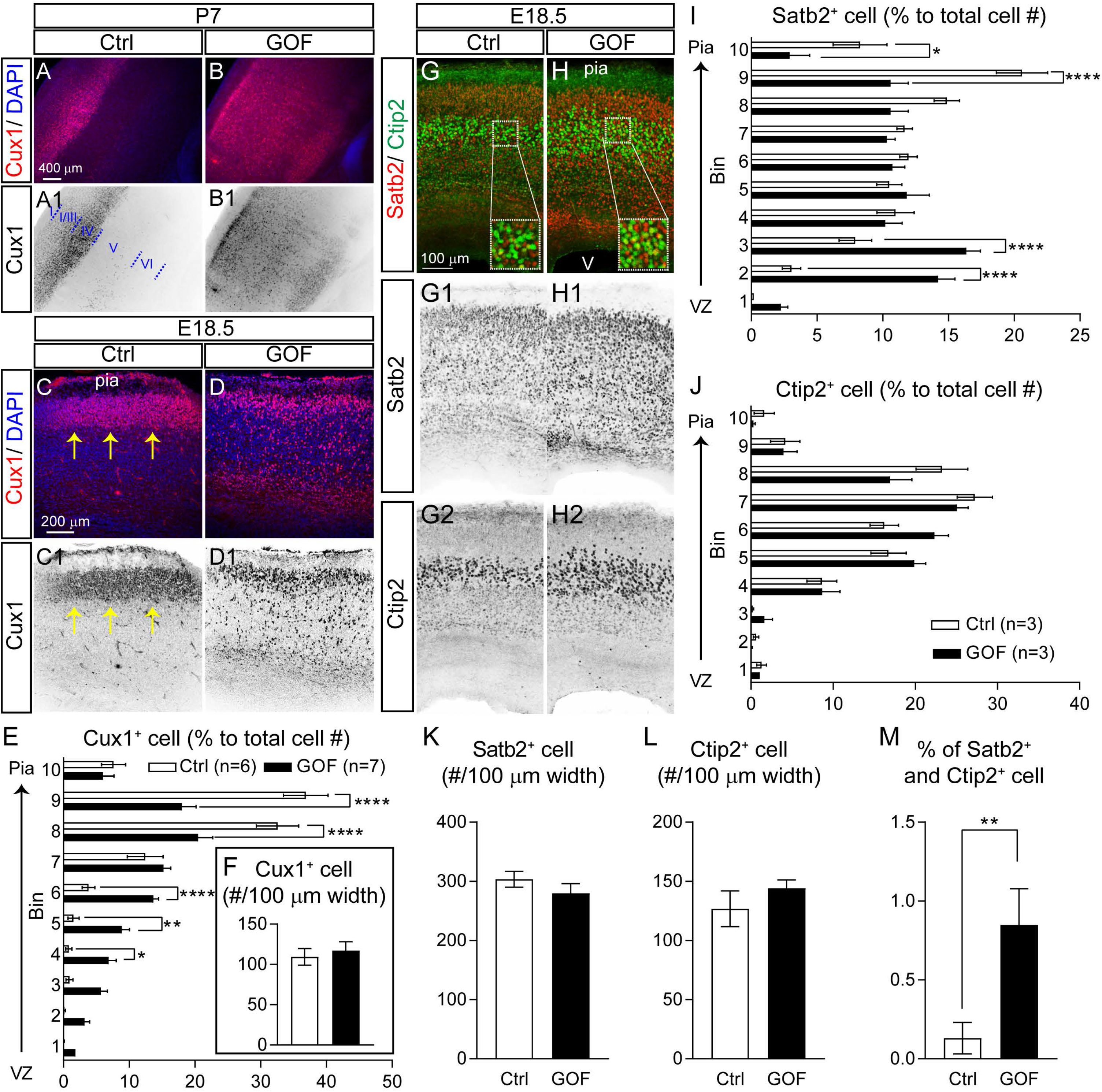
FGFR GOF in postmitotic neurons resulted in the misplacement of later-born principal neurons. (A, B) Example images of Cux1 staining with coronal sections prepared from P7 ctrl (control) and GOF S1 cortex. (C-D) Cux1 staining in E18.5 ctrl and GOF cortical plate. A1-D1 show the inverted Cux1 staining images. The yellow arrows indicate the cluster of Cux1^+^ cells. (E) The distribution of Cux1^+^ cells in E18.5 cortex alone different cortical depth. Two-Way ANOVA *post hoc* Bonferroni’s multiple-comparisons test. The statistical analysis (*) compared between ctrl and GOF for corresponding bins. *p<0.05; **p<0.01; ****p < 0.0001. (F) Summary of the total number of Cux1^+^ cells. Student’s-t test. (G, H) Example images for Satb2 and Ctip2 double staining with E18.5 Ctrl and GOF cortex. G1-2 and H1-2 show the inverted images of Satb2 and Ctip2 staining. (I, J) Summaries for the distributions of Satb2^+^ and Ctip2^+^ cells alone the cortical depth in the E18.5 cortex. Two-Way ANOVA *post hoc* Bonferroni’s multiple-comparisons test. (K, L) Summary graphs for the total numbers of Satb2^+^ and Ctip2^+^ cells. (M) Summary for the percentage of neurons co-expressing Satb2 and Ctip2. Student’s-t test. ** p<0.01. Pia, pial surface; V, ventricle. All the images shown were in the coronal plane.

### FGFR3 GOF in post-mitotic principal neurons results in a reduction of radial glia cells (RGCs) and a loss to their radial processes

Newly born glutamatergic neurons radially migrate towards the pial surface via radial glia assisted radial migration ^40,42^. The absence of cortical layers in GOF brains prompted us to examine the distribution of Cux1^+^ neurons and radial glia processes at embryonic day 15.5 (E15.5) when radial migration is active. Radial glia processes are derived from Pax6^+^ RGCs and can be visualized by staining with an antibody against a brain-specific member of the lipid-binding protein (BLBP)^43^. We found that some Cux1^+^ cells already migrated to the top of the cortical plate at E15.5 (Fig. 3A, A1) while the Cux1^+^ cells in the GOF brains remain stayed at the bottom of the cortical plate (Fig. 3B, B1). We also found that abundant BLBP^+^ radial processes were present in both rostral and caudal areas of the control embryonic cortex (Fig. 3C, C2, E, E2). However, the BLBP^+^ processes were barely detectable in GOF embryos (Fig. 3D2, F2). The presence of GFP in the cortical plate of the GOF embryos indicated Cre-mediated recombination occurred in post-mitotic neurons located in the cortical plate region (Fig. 3D, D1, F, F1). There was no GFP expression in control embryonic brain tissue (Fig. 3C, C1, E, E1). Neurons in the cortical plate tended to align in a columnar fashion, as a result of radial migration (Fig. 3C3, E3). In contrast, the arrangement of neurons in the GOF cortical plate appears rather disorganized (Fig. 3D3, F3). Results suggest that RGCs are affected by FGFR3 GOF in post-mitotic neurons.

**Figure 3.**
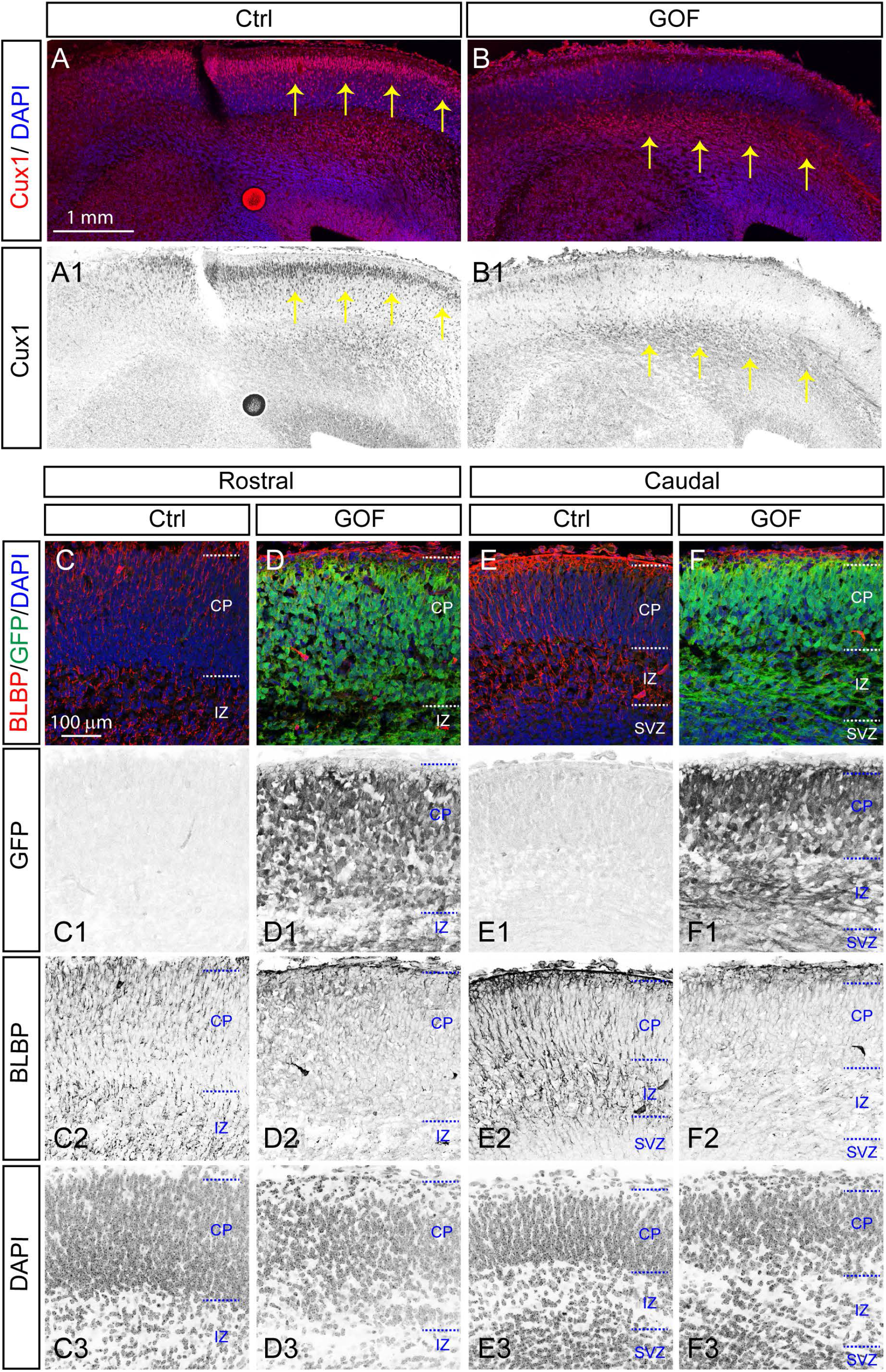
FGFR GOF in post-mitotic principal neurons results in a loss of radial processes. (A, B) Cux1 staining with coronal sections prepared from E15.5 ctrl (control) and GOF S1 cortex. A1 and B1 show the inverted Cux1 staining images. Yellow arrows indicate the Cux1^+^ cell. (C-F) Confocal images showed BLBP and GFP staining in rostral (C, D) and caudal (E, F) plane at E15.5 embryos. C1-F1, C2-F2, and C3-F3 show the inverted GFP, BLBP, and DAPI staining images. The GFP^+^ cells are in both cortical plate (CP) and intermediate zone (IZ) in GOF embryos. GFP expression in GOF embryos indicates the *FGFR3*^*K650E*^ allele is successfully activated at E15.5. Notably, BLBP^+^ processes are barely detected in the cortical plate and intermediate zone in GOF embryos. CP, cortical plate; IZ, intermediate zone; SVZ, subventricular zone.

RGCs not only assist newborn neurons to migrate in the appropriate direction; they also act as stem cells that divide in an increasingly asymmetric manner to self-renew and generate restricted intermediate progenitor cells and neurons ^12^. Thus, the finding that neuronal FGFR3 hyperfunction impaired RGC process formation suggested that the function of neural precursor cells may also be compromised. To examine whether neuroprogenitor cells, PAX6^+^ RGCs and Tbr2^+^ IPCs ^12–14,43–45^, were altered, we examined the number and distribution of PAX6^+^ and Tbr2^+^ cells. As expected, PAX6^+^ staining revealed RGCs arranged as a packed layer at the base of the VZ (Fig. 4A, A11) while Tbr2^+^ immunoreactivity was mainly found in the SVZ/VZ (Fig. 4C, C1). In contrast, the thickness of the GOF cortex was significantly less than control (p=0.0002; Fig. 4E) (Fig. 4B, B1, D, D1). Quantification of GOF cortex revealed a significant reduction of PAX6^+^ RGCs (p=0.0099; Fig. 4F), while the number of Tbr2^+^ IPCs was normal in GOF cortex (p=0.1571; Fig. 4G). In addition, there was no significant difference in the density of PAX6^+^ (Ctrl, 15254 ± 1479 number/ mm^2^ v.s. GOF, 14747 ± 993.4 number/ mm^2^; p=0.77) or Tbr2^+^ (Ctrl, 12181 ± 484.5 number/ mm^2^ v.s. GOF, 13861 ± 789.2 number/ mm^2^; p=0.0834) cells, likely due to the reduced cortical thickness in GOF brains.

**Figure 4.**
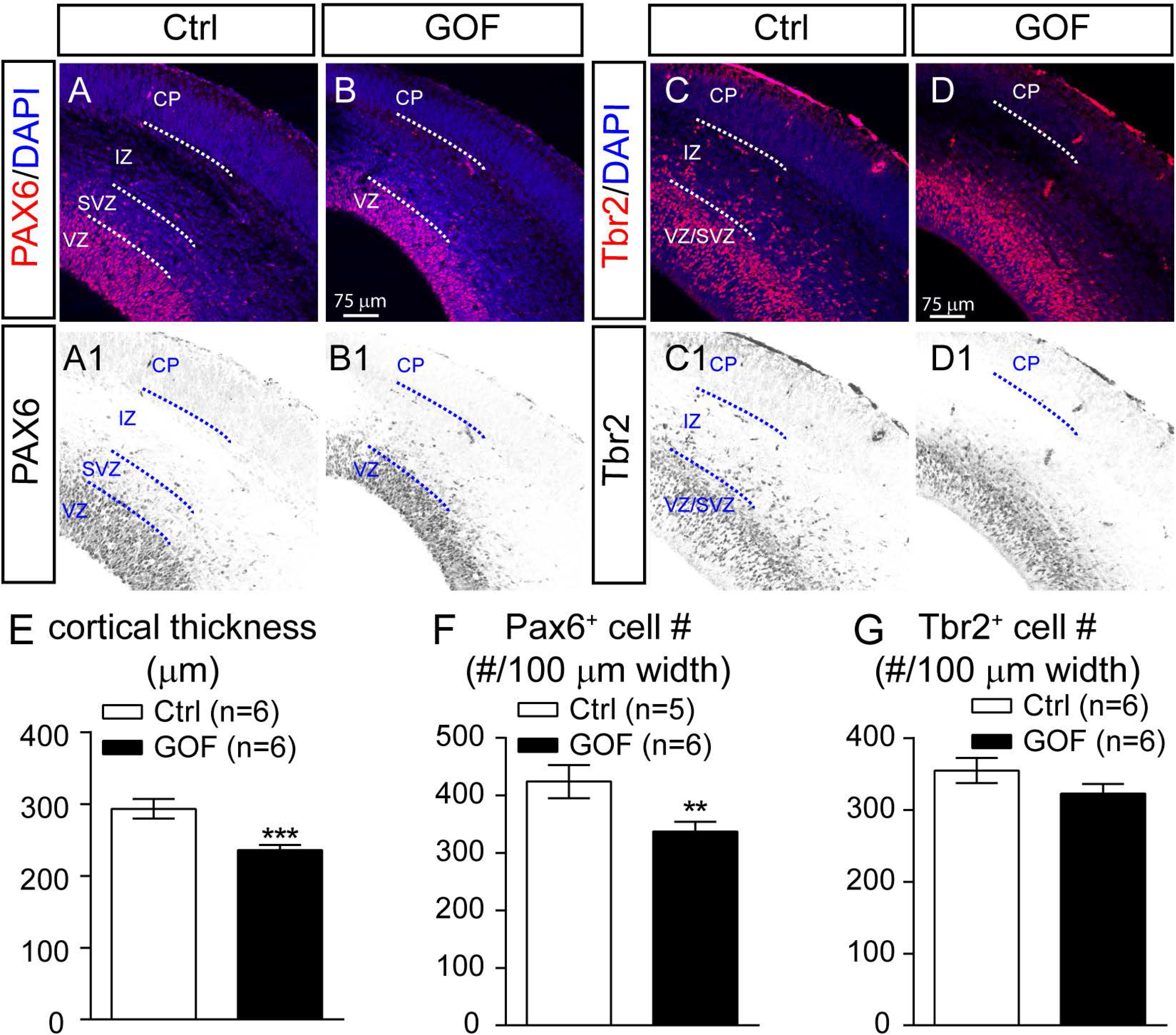
FGFR GOF in post-mitotic principal neurons results in a reduction of radial glia cells (RGCs). (A, B) Exemplary images of PAX6 and DAPI staining. A1 and B1 show the inverted images of PAX6 staining. (C, D) Exemplary images of Tbr2 and DAPI staining. C1 and D1 show the inverted images of Tbr2 staining. CP, cortical plate; IZ, intermediate zone; SVZ, subventricular zone. Summary for the measured cortical thickness (E), Pax6^+^ cell number (F), and Tbr2^+^ cell number (J). Student’s-t test. **p<0.01; ***p < 0.001.

To confirm that the loss of radial glia processes and reduced number of PAX6^+^ cells was due to non-cell autonomous influences of FGFR3 GOF from post-mitotic neurons instead of leaky Cre expression in PAX6^+^ cells from Nex-Cre line, we examined if Nex-Cre can induce recombination in PAX6^+^ cells. E15.5 embryos carrying one copy of the tdTomato reporter gene (Ai9, Rosa-CAG-LSL (stop)-tdTomato) and one copy of the Nex-Cre allele were generated and collected for Pax6/Tbr2 double immunostaining (Supplementary Fig. 3). tdTomato^+^ cells, reporting Cre-mediated recombination, are mainly localized at the cortical plate (Supplementary Fig. 3C, H) and their axons projected through IZ (Supplementary Fig. 3 C, E, G). No tdTomato cells were found to be Pax6^+^ or Tbr2^+^ (Supplementary Fig. 3D, F). Taken together, our results suggest that FGFR3 GOF in post-mitotic principal neurons imparts non-cell autonomous effects on RGCs.

### FGFR3 GOF in post-mitotic principal neurons decreases the number of neurons and glial cells and brain size

To determine whether the reduction of PAX6^+^ RGCs in the GOF embryonic cortex results in fewer cells in the postnatal brain, IdU was injected at E14.5 to label cells born at E14.5 and then the number of IdU^+^ cells in the S1 cortex was examined at P7 (Fig. 5A, B). We found that the number of IdU^+^ cells (p=0.0001; Fig. 5C), NeuN^+^ cells (neurons, p= 0.0061; Fig. 5D), and S100β^+^ cells (astrocytes, p<0.0001; Fig. 6E) were significantly decreased in the GOF cortex. However, there was no significant difference in the ratios of NeuN-IdU double-positive cells to total IdU^+^ cells between GOF and control cortex (p=0.4165; Fig. 5F). We also evaluated brain size by imaging the dorsal side of fixed adult (P145) GOF and control brains (Fig. 5G, H) and measured the projected dimensions. We found that the 2D projected length of olfactory bulb (p<0.0001; Fig. 5I) and midline (p=0.0004; Fig. 5J) of fixed brains as well as the projected cortical surface area (p<0.0001; Fig. 5K) were significantly less in the GOF mice compared to control. Our results suggest that FGFR3 GOF solely in post-mitotic neurons results in fewer RGCs and decreases the number of neurons and astrocytes in the cortex and decreases brain size.

**Figure 5.**
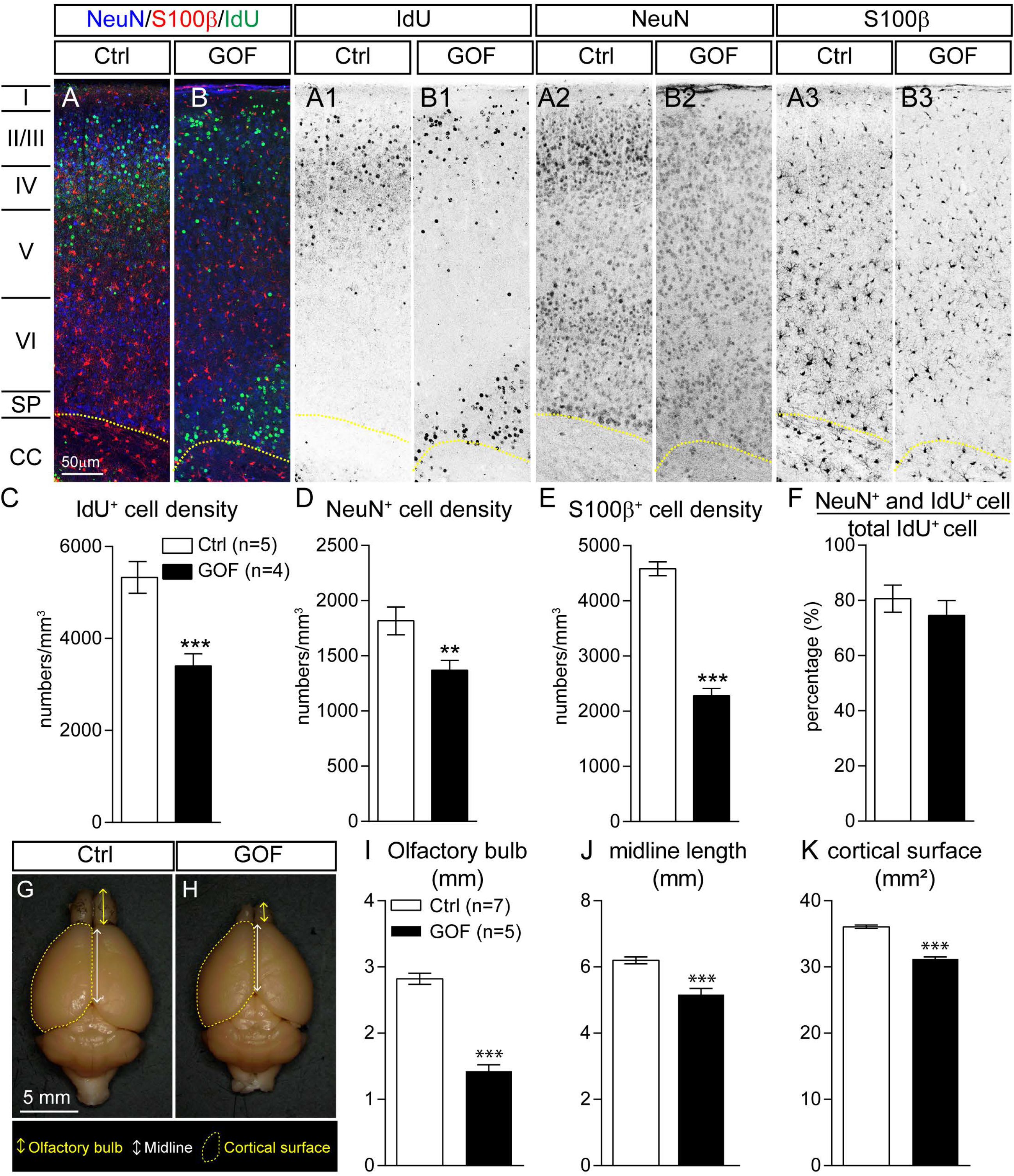
FGFR GOF in post-mitotic principal neurons decreases neurons, glia, and brain size. Time pregnant females at E14.5 were injected with IdU and their progenies were examined at P7. (A, B) Exemplary images show NeuN, S100β and IdU staining in P7 ctrl and GOF S1 cortex. A1-A4 and B1-B3 show the inverted images of IdU, NeuN, and S100β staining. Summaries for IdU^+^ (C), NeuN^+^ (D), S100β^+^ (E) cell densities as well as the percentage of NeuN^+^-IdU^+^ double-positive cells to IdU^+^ cells (F). (G, H). Representative images (dorsal view) of brains harvested from P121 Ctrl and GOF mice. The olfactory bulb length (I), midline length (J), and cortical surface area (K) in G and H were measured (Ctrl, n=7; GOF, n=5) with photos in 2-D. Student’s-t test. **p<0.01; ***p < 0.001; ****p < 0.0001. I-VI, cortical layers; SP, subplate. CC, corpus callosum.

**Figure 6.**
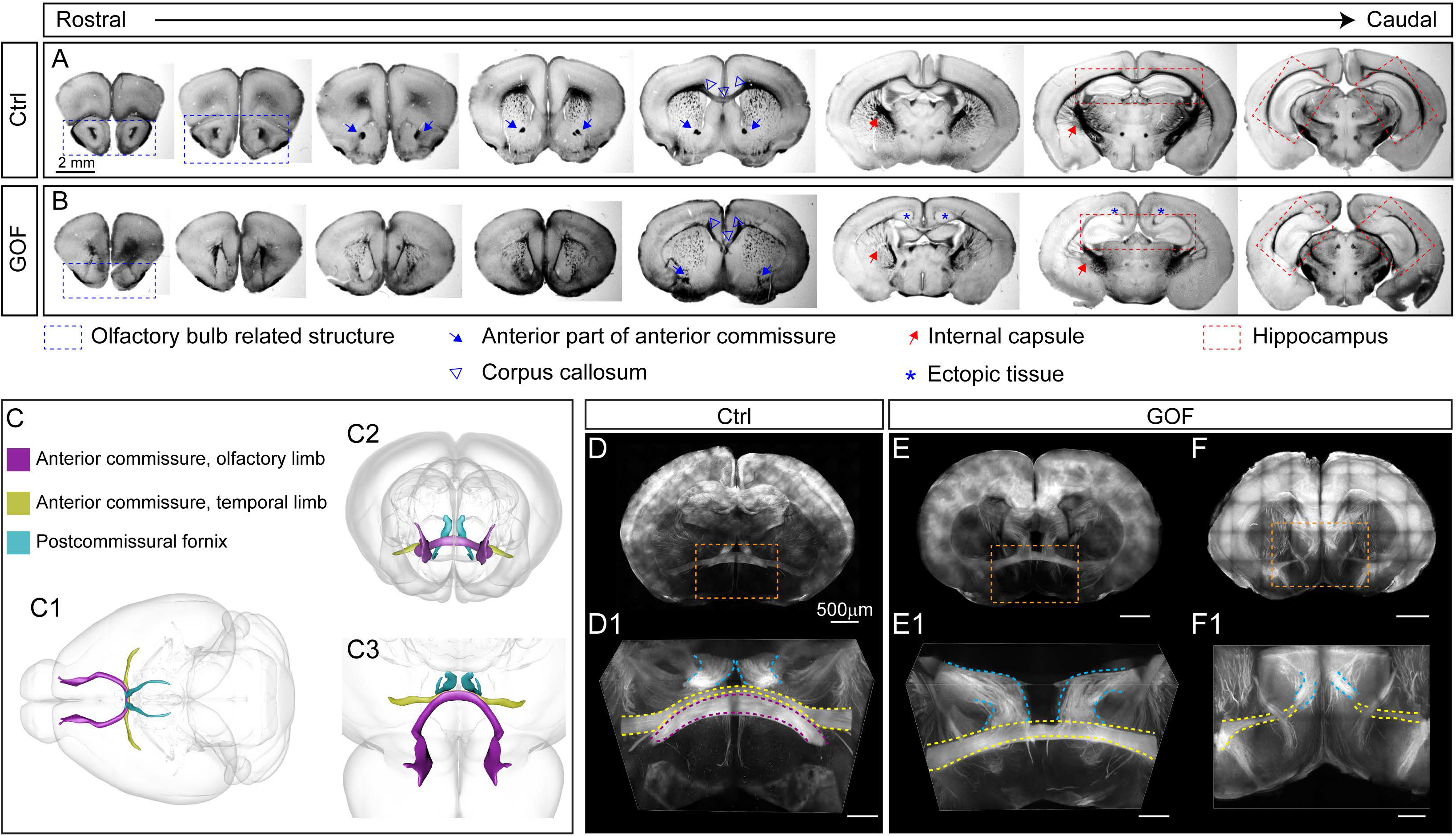
The anterior and posterior commissures are severely miswired in FGFR3 GOF mice. (A, B) Representative images show the anatomical features of coronal brain sections from rostral to caudal area in ctrl and GOF mice. The olfactory bulb related structures are marked with blue rectangles. The blue arrows indicate the anterior part of the anterior commissure. The red arrows indicate the internal capsule. The blue arrowheads indicate the corpus callosum. Blue stars mark the ectopic cell clusters in GOF mouse brains. The red rectangles mark hippocampal regions. (C) Scheme illustrating the axonal projections in 3-D. C1 shows the dorsal view. C2 shows the front view from the rostral side which is like the images in D-F. C3 represents the viewing orientation as the images in D1-F1. (D-F) The fluorescence images from ctrl and GOF coronal brain slices. D1, E1, and F1 show the enlarged views for the area indicated with orange rectangles in D-F. The arch-shaped olfactory limb of the anterior commissure is high-lighted with purple dashed lines. The temporal limb of the anterior commissure (labeled by dashed yellow lines) is located in front of the columns of postcommissural fornix (labeled by dashed blue lines). D, dorsal; V, ventral, A, anterior; P, posterior.

### FGFR3 GOF in post-mitotic principal neurons results in a severe axon miswiring

FGF-FGFRs have been shown to regulate axonal pathfinding during embryonic development ^46–49^, thus we further examined if FGFR3 GOF perturbed brain wiring. Serial brain sections were prepared and imaged under a dissecting bright-field microscope (Fig. 6). Several anatomical deficits in GOF mice were evident even at this gross level of evaluation. First, olfactory bulb related structures, including the anterior olfactory nucleus (AOD) and anterior olfactory nucleus external part (AOE), were smaller or not fully developed (Fig. 6B) in GOF mice. In control mice, the anterior part of the anterior commissure (ACA) passed through several coronal sections as a bundle passing through the ventral part of the brain (Fig. 6A). However, ACA fibers were not evident or were shifted more laterally in the GOF brain (Fig. 6B). The projection pattern of the internal capsule, which contains axonal projection fibers, including cortical thalamic, cortical striatal, and thalamocortical projections, was reduced and deformed (Fig. 6A, B). The thickness of the corpus callosum was greatly reduced (Fig. 6A, B). Ectopic cell clusters were also observed in GOF mouse brains (Fig. 6B). The overall shape of the hippocampi was quite round (Fig. 6A, B) in GOF mice.

The evident fiber tract abnormality at gross anatomical levels motivated us to further examine the impact of FGFR3 GOF on axonal trajectories that are originating from Nex-lineage neurons in 3-D. Thus, we generated the triple transgenic line with tdTomato reporter to label the axonal projections of Nex-lineage neurons with or without FGFR3 GOF. While the 2-D bright-field images reveal gross anatomical differences, examining brain tissue at 3-D level provides a more comprehensive understanding of axonal trajectories. With the combination of the ScaleS tissue clearing methods ^50^ and two-photon microscope imaging, we acquired image stacks of 1 mm-thick coronal brain sections to examine the axonal projection in 3 dimensions (Fig. 6D-F, Supplementary Fig. 4). We first examined the most anterior 1mm slices which containing parts of the olfactory and temporal limbs of the anterior commissure as well as the postcommissural fornix (Fig. 6C). We found that the olfactory limb of the anterior commissure showed an “arch shape” (Fig. 6D1), yet, this structure is missing in GOF mice (Fig. 7E1, F1). The temporal limb of the anterior commissure, located in front of the postcommissural fornix, was observed in all control but only in “some” GOF mice (Fig. 6D1, E1, Supplementary Fig. 4). Intriguingly, some GOF mice had apparent aberrant temporal limb of the anterior commissure that were not crossing midline and abnormal trajectories (Fig. 6F1, Supplementary Fig. 4). In summary, the posterior portion of the olfactory limb of the anterior commissure appears to be heterogeneously altered in GOF mice.

**Figure 7.**
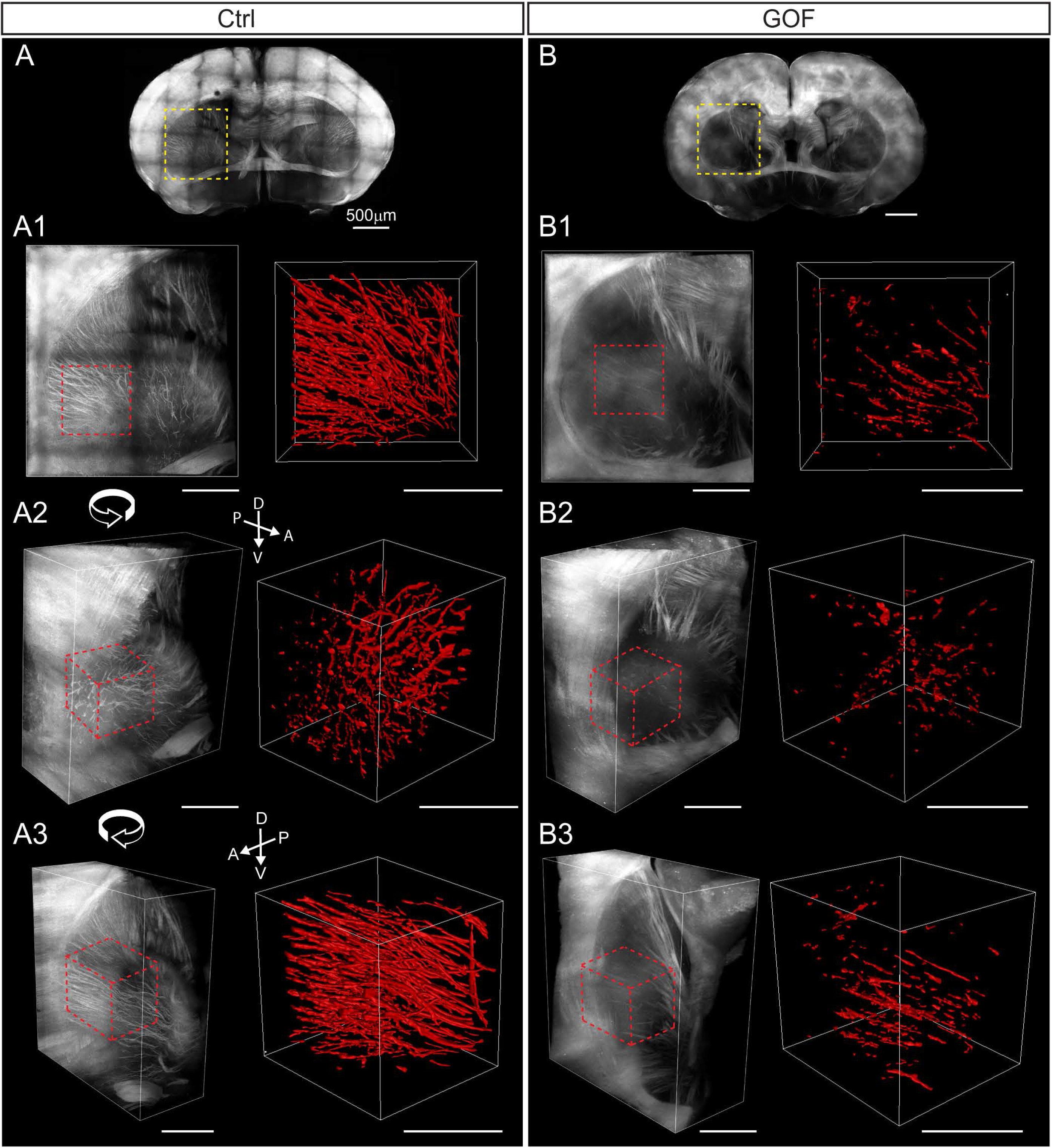
FGFR3 GOF mice have fewer axonal projections in the striatum. (A, B) Overview of coronal brain slices from control (Ctrl) and GOF mice. A1-A3 and B1-B3 show the zoomed-in view with different rotation angles in the striatum (yellow rectangle in A and B). The region analyzed in Imaris is marked with dashed red lines. The right panel in A1-A3 and B1-B3 shows the 3-D rendering of axonal fibers in the striatum (dotted-line delimited area in the left panel). D, dorsal; V, ventral, A, anterior; P, posterior.

TdTomato^+^ axonal projections reveal cortico-striatal and cortico-thalamic projections in the striatum in control mice (Fig. 7). These tdTomato^+^ long-range cortical axons were greatly reduced in GOF striatum (Fig. 7, Supplementary Fig. 5). We selected a portion of the striatum located in a similar position and applied Imaris surface rendering function to reconstruct the tdTomato^+^ axonal tracts in both control and GOF brains (Fig. 7A1-A3, B1-B3). The quantification reveals that tdTomato^+^ axonal projections in the selected striatal area occupied a significantly smaller volume in GOF than control mice (Ctrl, n=5, 17.7 ± 3.096 ×10^6^ μm^3^ v.s. GOF, n=4, 2.595 ± 0.5197 ×10^6^ μm^3^; p=0.0037). Taken together, the number of Nex-lineage axonal projections present in the striatal area is significantly reduced in GOF mice.

The postcommissural fornix, the major output from the hippocampal subiculum, innervates the anterior hypothalamus through the medial-corticohypothalamic tract and terminates at the posterior hypothalamus in the mammillary body ^51,52^. We utilized the same approach described above to quantitatively characterize the postcommissural fornix in ctrl and GOF brains (Fig. 8, Supplementary Fig. 6). The bundles of the postcommissural fornix in control brains are tightly packed and exhibit a prolate ellipsoid, or a “cigar-shaped” (Fig. 8B1. B3). However, these the postcommissural fornix bundles in GOF brains are loosely packed and resemble an oblate ellipsoid, or a “disc-shaped” structure (Fig. 8C1, C3). These bundles in GOF mice have a significantly smaller value of prolate ellipticity (p=0.0102; Fig. 8D) and a significantly higher value of oblate ellipticity (p<0.0001; Fig. 8E) compared to control bundles. There is no significant difference in total volume occupied by tdTomato^+^ axonal projections (p=0.1407, Fig. 8F) between the two lines. These data suggest similar numbers of axons between ctrl and GOF postcommissural fornix. To determine if the shape change is caused by axonal fasciculation deficits by FGFR3 GOF, we analyzed the diameter of the bundles in 4 horizontal planes extracted from the 3D volume for both ctrl and GOF brains (Fig. 8B4, C4). A significant increase in the diameter of the postcommissural fornix bundles found in GOF compared to control mice (p<0.0001, Fig. 8G) indicate FGFR3 GOF perturbs axonal fasciculation of the postcommissural fornix.

**Figure 8.**
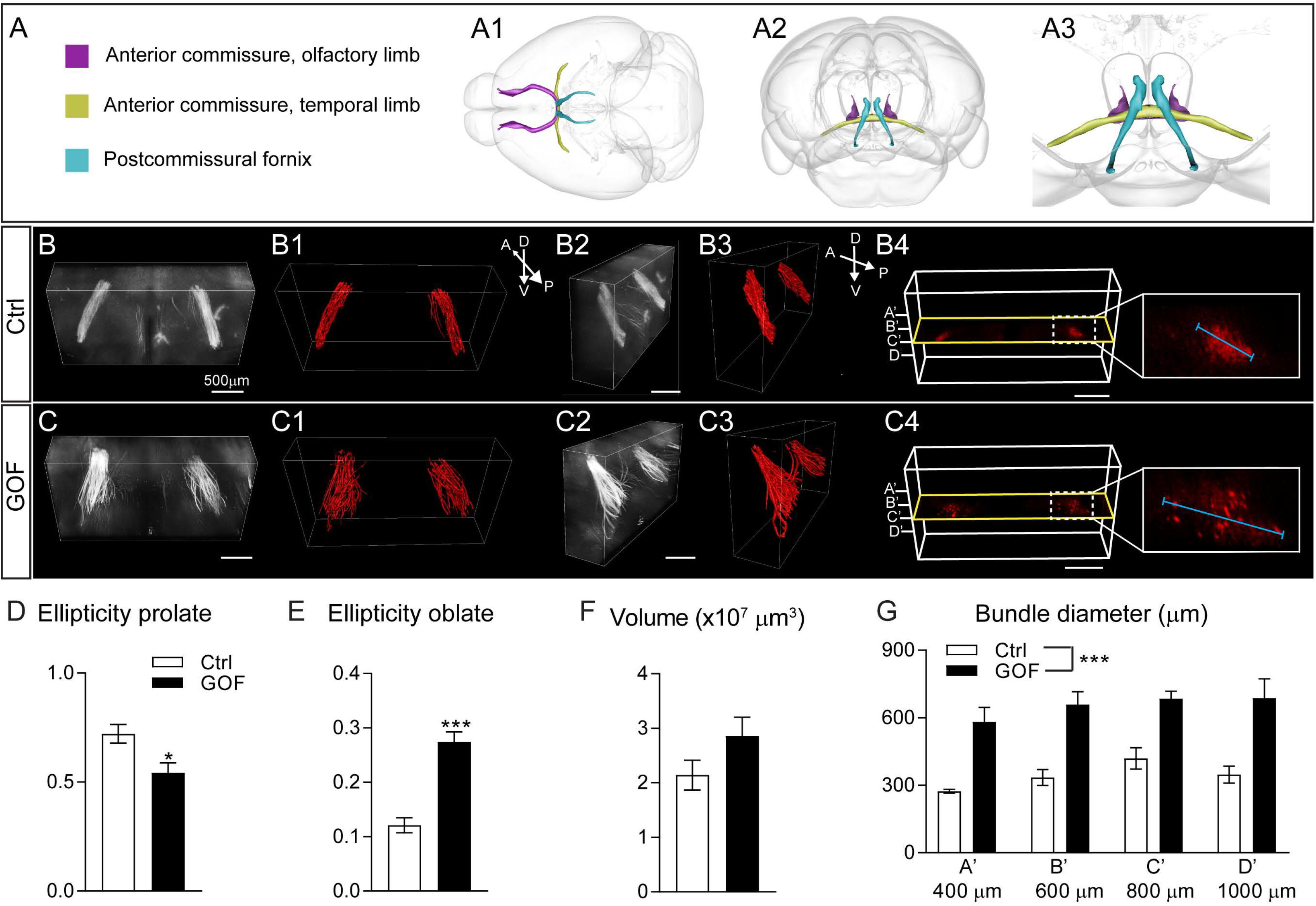
FGFR3 GOF disrupts axonal fasciculation of the postcommissural fornix. (A) Scheme illustrating the axonal projections in 3-D. A1 shows the dorsal view. A2 shows the front view from the roster side. A3 represents the enlarged view in A2 (B, C) The front views from the roster side of the postcommissural fornix bundles from ctrl and GOF mice. B2 and C2 show the side view of B and C. B1, B3, C1, and C3 show the Imaris 3-D surface rendering images. (B4, C4) Scheme illustrating the image planes used to measure the diameter of the postcommissural fornix bundle. 2-D images from four planes (A’, B’, C’, D’ at 400μm, 600μm, 800μm, and 1000μm) were extracted and the longest diameter of the axonal bundle in each plane was measured. Summaries for prolate ellipticity (D), oblate ellipticity (E), total volume occupied by postcommissural fornix projections (F). Student’s-t test. (G) Summary of the diameters of postcommissural fornix bundles at four different planes indicated in B4 and C4. Statistical analysis: Two-Way ANOVA *post hoc* Bonferroni’s multiple-comparisons test. D, dorsal; V, ventral, A, anterior; P, posterior.

### Unbiased transcriptomic analysis reveals putative cellular mechanisms on how FGFR GOF impacts RGCs, neuronal differentiation, and axonal pathfinding

To further explore the putative molecular mechanisms on how FGFR3 GOF impacted cortical development, RNA-seq was conducted with RNA prepared from E15.5 control and GOF embryonic cortices as the mislocalization of Cux1^+^ cells was already evident at E15.5. RNA-seq studies found that FGFR3 GOF in glutamatergic neurons resulted in 537 genes upregulated and 596 genes down regulated (FDR < 0.05; Fig. 9A, Supplementary Dataset 1). Based on the RNA-seq data, mRNA levels of RGC markers such as *Pax6*, *Fabp7* (encoded brain-specific member of the lipid-binding protein, BLBP, which is located in RGC processes), and *Nestin* as well as the IPC specific gene marker *Enomes* (Tbr2, IPCs) were decreased in GOF cortices (Fig. 9B). Additionally, mRNA levels of several neuronal markers such as *Tbr1*, *Cux1*, *Satb1*, *Satb2* did not differ between control and GOF cortices (Fig. 9B). An exception to this was that the expression of *Bcl11b* (Ctip2), a marker of deep layer cortical neurons was significantly upregulated (Fig. B).

**Figure 9.**
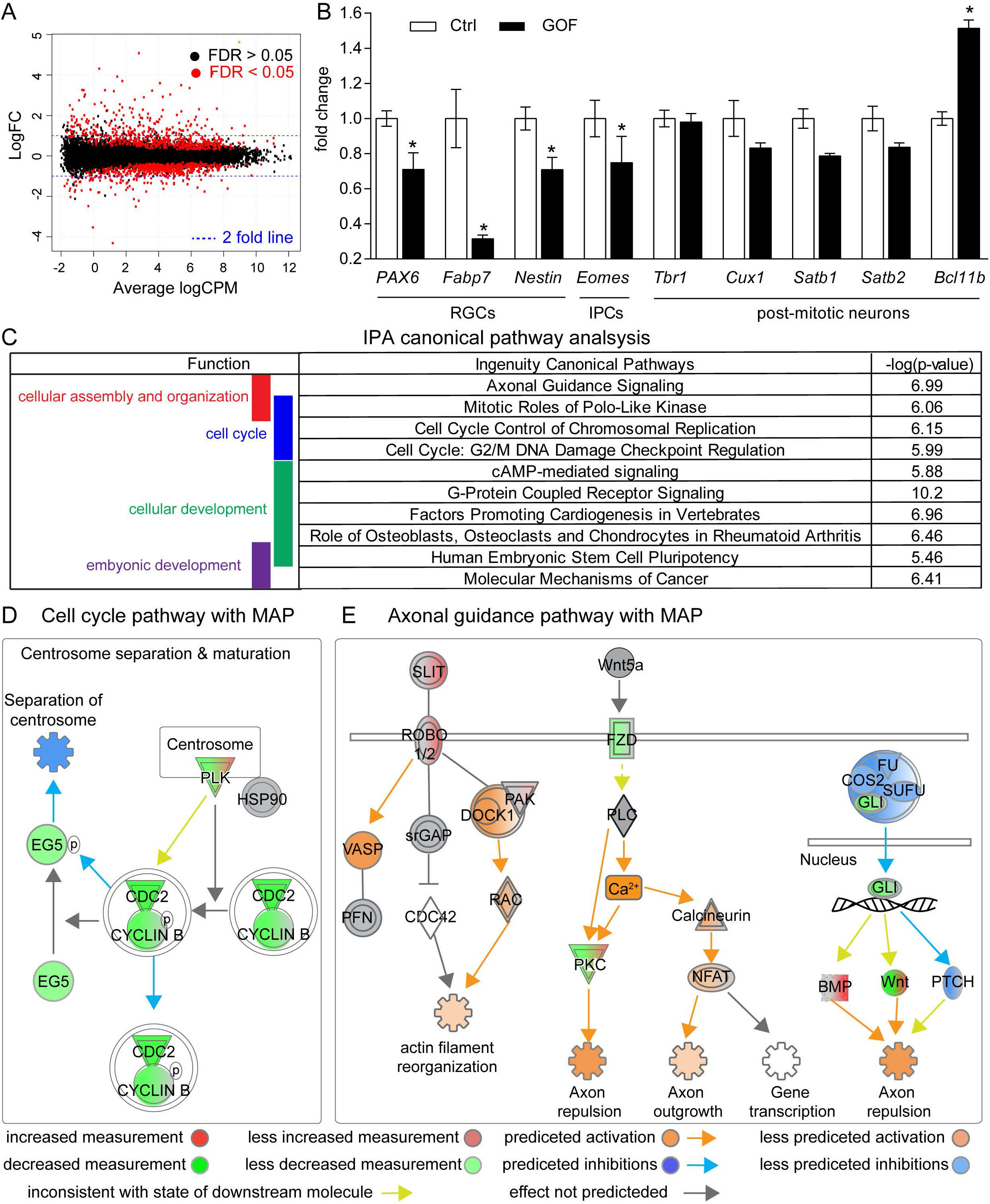
IPA analysis shows strong suppression in cell cycle progression genes and dysregulation in axon guidance pathways in E15.5 GOF embryos. (A) The smear plot demonstrates the Log fold change (FC) against average log counts per million reads (CPM). The black and red dots represent genes with FDR > 0.05 and FDR < 0.05, respectively. Blue lines indicate the 2-fold change lines. (B) Transcriptional changes of genes associated with RGCs (*Pax6*, *Fabp7, Nestin*), IPCs (*Eomes*/Tbr2), and postmitotic neurons (*Tbr1*, *Cux1*, *Satb1*, *Satb2*, *Bcl11b*/ctip2) summarized from RNAseq data. (C) The top 10 significant canonical pathways (based on p-value) identified by IPA analysis classified based on physiological function. These pathways are involved in regulating cellular assembly and organization, cell cycle, cellular development, and embryonic development (showed in the left panel). (D, E) Schematic diagrams show the partial pathways related to cell cycle and axonal guidance with the molecule activation prediction (MAP) analysis. Molecules are indicated by standard abbreviations. Relative changes in gene expression are depicted by graduated shades of color coding: red, up; green, down; white, no change or not applicable. MAP analysis in IPA predicts the activation status of each pathway component based on the transcriptional state of the relevant genes. Relative activation and inhibition status are depicted by graduated shades of color coding: orange, activation; blue, inhibition; yellow, inconsistent with the state of the downstream molecule; grey, effect not predicted.

We further conducted Pathway analysis with Ingenuity Pathway Analysis (IPA; Qiagen) to aid in the interpretation of downstream biological pathways affected by FGFR3 GOF. Pathway analysis found 185 canonical pathways that had z-scores either considerably greater or considerably less than 0 and for which the up-down-regulation was statistically significant (Supplementary Dataset 2), including expected FGF signaling and ERK/MAPK signaling. When the top 10 significant canonical pathways were broadly classified based on physiological function as described in the IPA database, pathways involved in regulating cellular assembly and organization, cell cycle, cellular development, and embryonic development were prominent (Fig. 9C).

The decrease of RGC number and robust axon miswiring prompted us to explore further the cell cycle-regulated pathways and the axonal guidance signaling pathway in-depth. We found pathways that regulate cell cycle were specifically related to mitosis most affected (Fig. 9D). To determine the activation state of the pathways identified from IPA analysis, we performed a molecule activation prediction (MAP) analysis in IPA. This bioinformatics tool predicted the activation states of each pathway component based on the transcriptional state of the relevant genes. MAP analysis showed that the centrosome separation pathway was inhibited (Fig. 9D). In the pathway of centrosome separation and maturation, several key downstream effectors proteins such as kinesin family member 11 (EG5), cell division control protein 2 (cdc2), and cyclin B were downregulated in FGFR3 GOF embryos (Fig. 9D). In the axonal guidance signaling pathway, a total of 50 genes were altered in FGFR3 GOF embryos including *Slit2*, *Slit3*, *Robo1, Fzd1, Fzd2, Fzd8, Gli3, and Wnt* (Fig. 9E). MAP analysis predicted that several events related to axonal outgrowth such as actin filament reorganization, axon repulsion, and axon outgrowth are activated (Fig. 9E). Taken together, the transcriptomic profile suggests that FGFR3 GOF in post-mitotic glutamatergic neurons exerts non-cell autonomous influences on cell cycle progression of NPCs’s and a cell-autonomous impact on neuronal pathfinding.

## Discussion

In this study, we test specifically how FGFR3 GOF in the post-mitotic neurons impact brain development. Using the GOF approach, we extend previous studies to reveal that FGFR signaling hyperactivation exerts significant adverse effects on cortical development. FGFR3 GOF in post-mitotic neurons is sufficient to impart non-cell autonomous effect on RGCs, reducing RGCs numbers and function, and exert cell-autonomous effect to affect axonal guidance, resulting in overall abnormal cortical lamination and axon miswiring. The results reported here substantially extend our understanding of neuronal FGFR action in CNS.

### FGFR3 GOF in NEX-lineage neurons leads to aberrant cortical and hippocampal lamination

Radial migration guided by RGCs is a critical cellular process in cortical lamination ^42,53,54^. Hippocampal neurons located in the CA1 region also require radial glial fibers for their appropriate migration ^55^. The greatly reduced RGCs’ radial processes in the cortical plate of GOF embryonic brains (Fig. 3C-F) suggest the detective radial migration account, at least in part, for the lamination defects. The observation that migration defects mainly affects upper-layer neurons (Cux1^+^ and Satb2^+^) rather than in deeper layer neurons (Ctip2^+^) neurons further supports this notion (Fig. 2). However, our studies cannot rule out the possibility that FGFR3 GOF in neurons perturbs their somal translocation ability during the migration process, with subsequent loss of RGC radial processes. Interestingly, deleting GSK3 using NEX-Cre resulted in radial migration and lamination phenotypes ^56^, similar to the phenotypes we observed in GOF brains. In Morgan-Smith’s study, they showed that GSK3 deletion delays the multipolar to bipolar transition for migrating cortical neurons. It has been shown that FGFR signaling activates Akt signaling ^57^, which subsequently inhibits GSK3 activity by phosphorylating GSK3. Thus, it is plausible that FGFR3^K650E^ expression in cortical neurons results in reduced GSK3 signaling and alters neuronal migration.

### FGFR3 GOF in NEX-lineage neurons perturbs RGC cells and reduces brain size

The formation of neural circuits within the central nervous system depends upon the precise spatially and temporally coordinated generation of distinct classes of neurons and glia from RGCs and IPCs ^23,58^. Our finding that PAX6^+^ RGCs are decreased in GOF embryonic cortex (Fig. 4) likely accounts for the reduced number of neurons and astrocytes in postnatal GOF brains as well as their smaller brain size (Fig. 5). IPA analysis of the transcriptomic profile of the E15.5 GOF embryonic cortex identifies that canonical pathways related to cell cycle progression are altered (Fig. 9). Specifically, MAP analysis predicted that the downregulation of polo-like kinase (PLK1) may be the upstream protein to suppress cell cycle progression (Fig. 9D, Supplementary Dataset 2). PLK1, a serine/threonine-protein kinase, has been recognized as a key regulator of mitosis, meiosis, and cytokinesis ^59,60^ and is expressed in VZ/SVZ^61^. It remains to be experimentally validated for the pathways identified by bioinformatics with future experiments and if FGFR3 GOF impairs the proliferation of RGCs.

We postulate that FGFR3 GOF in post-mitotic cortical neurons exerts non-cell autonomous influences on RGCs, resulting in the reduction of RGCs and the decrease in cell cycle progression gene expression in GOF mice. When an Ai9 Cre reporter mouse (loxP-flanked STOP cassette preventing transcription of a CAG promoter-driven tdTomato) was crossed to our colony’s NEX-Cre line to determine the sites of NEX-Cre mediated recombination, we found that tdTomato^+^ cells did not overlap with Pax6^+^ RGCs at E15.5 (Supplementary Fig. 3), consistent with previous findings ^29,62^. GFP derived from the FGFR3 floxed construct also highlighted the upper cortical plate in the VZ/SVZ area as the site of Cre-mediated recombination (Fig. 3D, F). Thus, the data presented here indicate that aberrant FGFR signaling in post-mitotic neurons impairs RGC functions. Such cross-talk between post-mitotic neurons and RGCs had been shown for cardiotrophin-1 ^63^, FGF18 ^64^, Sip1-NTF3 ^65^, FGF9 ^66^, and BDNF-BMP7 ^67^ signaling. It remains to be determined whether any of these signaling pathways are altered by FGFR3 GOF.

### FGFR3 GOF in NEX-lineage neurons leads miswiring of several long-range axonal tracts

Successful axon navigation depends on the competence of the growing tip of the axon to receive and integrate information provided by multiple spatially organized molecular cues arranged along the axon’s trajectory. Utilizing ScaleS clearing brain methodology, we visualized several miss-routed axonal tracts caused by FGFR3 GOF in cortical glutamatergic neurons. These tracts include anterior commissure, cortical striatal projection, and post-commissural fornix (Fig. 6-8). Interestingly, the miswiring phenomenon is heterogeneous in individual FGFR3 GOF mice (Fig. 6, Supplementary Fig. 4). RNA-seq data from E15.5 brain tissue identified significant dysregulation in axonal guidance signaling pathways including Slit-Robo, Wnt-Frizzled, and Gli pathway (Fig. 9E). In line with our observations, it has been shown that FGFRs regulate axon guidance cues such as slit and semaphorin3A to control axonal pathfinding ^46,49^. Slit-Robo signals regulate the pathfinding of multiple axonal pathways, including corticofugal, thalamocortical, and callosal connection ^68–71^. Frizzled-3, one of the members in Wnt-Frizzled pathway, is required for the development of multiple axon tracts including anterior commissure, corticofugal axons, thalamocortical axons, and corpus callosum in the mouse CNS ^72–75^. When frizzled-3 was removed in the neocortex (Emx1-IRES-Cre), the posterior part of the anterior commissure was completely missing and axons with aberrant trajectories appeared in the external capsule ^72^.

The molecular identity of neurons also determines the neuronal subtype and influence axonal wiring patterns. RNAseq studies revealed that the mRNA expression level of ctip2 (*Bcl11b*) was significantly upregulated in the E15.5 FGFR3 GOF brain (Fig. 9B). While CTIP2 immunostaining also revealed increased abundance at the protein level, without changing the total number of ctip2^+^ neurons (Fig. 2L). We also found that the percentage of satb2^+^/ctip2^+^ neurons was significantly increased in FGFR3 GOF cortices (Fig. 3M). It has been shown that Ctp2^+^/Satb2^+^ neurons project to callosal and sub-cerebral targets ^76^. Furthermore, ectopic expression of ctip2 in layer 2/3 neurons is sufficient to alter the axonal targeting of corticocortical projection neurons and cause them to project to subcortical targets ^77^. Thus, it is also possible that the miswiring phenomenon we observed in FGFR3 GOF mice is partially due to increased ctip2 expression in FGFR3 GOF neurons. Ctip2 expression is directly regulated by the satb2 transcriptional factor, which acts as a transcriptional repressor to inhibit ctip2 promoter activity ^78,79^. A recent study further shows that the transcriptional adaptor, Lmo4, inhibits Ctip2 by competing with Satb2 for Hdac1 binding ^76^. Interestingly, we found that *Lmo4* mRNA is significantly upregulated (FDR=0.0476, Fig. 10-1) in FGFR3 GOF brain tissue. Taken together, our data suggest that FGFR3 GOF may increase the *ctip2* expression level via upregulating Lmo4.

### The impact of FGFR3 GOF varies at different stages of cortical development

*In-situ* hybridization data reveal that FGFR1-3 mRNA levels are abundant in RGCs and IPCs during cortical development and but relatively low in the cortical plate, presumably post-mitotic neurons ^80^ (http://developingmouse.brain-map.org/). Despite the low expression level, the recent RNA-seq data demonstrated that postmitotic neurons also express FGFRs ^81,82^. These findings suggest that the downregulation of FGFR3 signaling may be important for maintaining the balance of neurogenesis and neuronal differentiation during cortical development. Several FGFR3 *de novo* mutations were identified in TD patients, including R248C, S249C, K650M, K650E ^83–87^. Patients with these mutations have intellectual disability, seizures, and cortical malformations ^83,84,88^. Lin et al. provided the first evidence that neural FGFR3^K650E^ expression directly impacts brain development ^19^. They generated CNS and cartilage-specific *Fgfr3*^*K644E*^ (corresponding to K650E in humans) mouse models with Nestin-Cre and Col2a1-Cre, respectively, to express *Fgfr3*^*K644E*^ (corresponding to K650E in humans) under regulation by its endogenous enhancer/promoter. The larger brain and cortical thickness in the Nestin-Cre *Fgfr3*^*K644E*^ mouse were attributed to increased proliferation and reduced apoptosis of progenitor cells ^20,89^. Ectopic neurons accumulating in the dentate gyrus were also observed ^19^. In contrast to their studies, the GOF mouse generated in this study by overexpressing *FGFR3*^*K650E*^ in NEX-lineage neurons exhibited a severe cortical lamination defect (Fig. 1, 2) and abnormal axonal tracts (Fig. 6-8, Supplementary Fig. 4-6) as having been observed in TD human patients but have not been reported in previous TD animal models. Our observation that the abnormal cortical lamination and hippocampal patterning is apparent only when FGFR3 is overexpressed in post-mitotic neurons, but not postnatally (Supplementary Fig. 2), further underscores the importance of a tightly regulated developmental window. In summary, our data suggest FGFR3 GOF in post-mitotic glutamatergic neurons may contribute to brain anatomical changes in TD patients. However, this hypothesis remains to be tested in the animal models where FGFR3 expression is controlled by its endogenous promoter. Nevertheless, the different stages of onset for *FGFR3*^*K650E*^ expression, for example in progenitor cells vs post-mitotic neurons, clearly impacts brain structures very differently.

## Materials and Methods

### Experimental Design and Animals

The mutant *FGFR3* allele (CAG-flox-stop-flox-*FGFR3*^*K650E*^-IRES-eGFP) and NEX-Cre transgenic mice were described previously ^29,30^. Specifically, a K650E mutation in *FGFR3* was identified in one form of neonatal lethal dwarfism, thanatophoric dysplasia II (TDII, OMIM 187601). This mutation is located in the activation loop of the kinase domain and causes constitutive activation of FGFR3 ^90,91^. The NEX-Cre allows Cre-mediated recombination to occur in post-mitotic cortical and hippocampal glutamatergic neurons from embryonic day 11.5 (E11.5) ^29^. The mixed genetic background from multiple crosses may result in a heterogeneous phenotype. To minimize this effect, littermate controls were used for all the experiments and processed simultaneously with hyperfunction samples. Mice carrying only NEX-Cre or flox *FGFR3*^*K650E*^ allele were used as controls. Heterozygous offspring were also crossed to Ai9 reporter mice (Rosa-CAG-LSL-tdTomato-WPRE; JAX Lab # 007909) to label axon projections of all NEX-lineage principal neurons. Both male and female mice were used in this study. All mice were housed in standard conditions with food and water provided ad libitum and maintained on a 12 hr dark/light cycle. All animal procedures were performed under Indiana University Bloomington Institutional Animal Care and Use Committee guidelines.

### Genotyping

Ear or tail lysates were prepared by immersing tissue pieces in digestion buffer (50 mM KCl, 10 mM Tris-HCl, 0.1% Triton X-100, 0.1 mg/ml proteinase K, pH 9.0), vortexing gently, and then incubating for hrs at 60°C to lyse the tissues. These lysates were then heated to 94°C for 10 minutes to denature the proteinase K (Thermo Scientific, Rockford, IL, USA), and centrifuged at 16,100g for 15 minutes. The supernatants were used as DNA templates for polymerase chain reactions (PCRs, EconoTaq Plus Green 2X mater mix, Lucigen, Middleton, WI, USA). The genotyping primers were as described previously^29,30,92^

### Chemicals and Antibodies

All reagents and chemicals were purchased from Sigma (St. Louis, MO, USA) unless otherwise stated. D(-)-Sorbitol, glycerol, and urea were purchased from Wako Chemicals USA Inc. (Richmond, VA, USA). Rabbit anti-phosphorylated FGFR substrate 2α (FRS2α, Tyr196, Cat# 3864, RRID: AB_2106222), rabbit anti-phosphorylated extracellular signal-regulated kinase 1/2 (ERK1/2, Thr202/Tyr204, Cat#9101; RRID: AB_331646), and mouse anti-ERK1/2 (Cat# 9107; RRID: AB_2235073) antibodies were purchased from Cell Signaling (Beverly, MA, USA). Rabbit anti-FRS2α (Cat# SC-8318; RRID: AB_2106228) and rabbit anti-Cux1 (Cat# SC-13024; RRID: AB_2261231) antibodies were purchased from Santa Cruz Biotechnology (Santa Cruz, CA, USA). Chicken anti-green fluorescent protein (GFP, Cat#GFP-1020, RRID: AB_10000240) antibody was purchased from Aves Labs (Tigard, Oregon, USA). Rabbit anti S100β (Cat# Z0311; RRID: AB_10013383) antibody was purchased from DAKO (Produktionsvej, Glostrup, Denmark). Guinea pig anti-vesicular glutamate transporter 2 (VGluT2) (Cat#2251-I; RRID: AB_2665454) and Rabbit anti-NeuN (Cat# MAB377; RRID: AB_2298772) was purchased from Millipore (Temecula, CA, USA). Rabbit anti-PAX6 (Cat#901301; RRID: AB_2565003) was purchased from BioLegend (San Diego, CA, USA). Rabbit anti-brain lipid-binding protein (BLBP, Cat# ab32423; RRID: AB_880078) and rabbit anti-Tbr2/Eomes (Cat# ab23345; RRID: AB_778267) antibodies were purchased from Abcam (Cambridge, MA, USA). Rat anti-BrdU antibody (Cat# OBT0030G; RRID: AB_609567; which recognized the BrdU derivative IdU) was purchased from Accurate Chemical (Westbury, NY, USA). All Alexa Fluor series conjugated secondary antibodies were purchased from Invitrogen (Grand Island, NY, USA). Secondary antibodies used for Western Blotting were purchased from LI-COR Biosciences (Lincoln, NE, USA).

### Western Blotting

For total protein extraction, brain tissue was lysed and homogenized with a modified radioimmunoprecipitation assay buffer [50 mM of Tris-base (pH 7.4), 50mM of NaCl, 1% Triton X-100, 0.1% sodium dodecyl sulphate, 1 mM of EDTA, 1% Na-deoxycholate, 1 mM of phenylmethylsulfonyl fluoride, 1 μg/mL of leupeptin, 1 μg/mL of aprotinin, 1 mM of Na_3_VO_4_, and 1 mM of NaF]. The supernatant solution was collected using centrifugation at 12000×g for 15 minutes at 4°C. Protein concentration was measured using a Bradford assay (Bio-Rad, Hercules, CA, USA). Proteins were separated on a 10% SDS-polyacrylamide gel and then transferred to a nitrocellulose membrane (BioRad, Boston, MA, USA). The membrane was incubated with the appropriate primary antibody and then incubated with a species-appropriate secondary antibody. Western blot images were acquired by LI-COR Odyssey scanner and software (LI-COR Biosciences, Lincoln, NE USA) and quantified with NIH ImageJ software.

### Immunostaining

Mouse brain tissues were prepared by intracardiac perfusion with phosphate-buffered saline (PBS) followed by 4% paraformaldehyde (PFA) prepared in PBS. Fixed brains were sectioned into 100 μm thick sections in the coronal plane using a Leica VT-1000 vibrating microtome (Leica Microsystems). Embryonic brain tissues were dissected at specific post-gestation dates, and post-fixed with 4% PFA for 2 hours at 4°C, followed by cryoprotection in 30% sucrose and embedding in optimal cutting temperature compound. Fixed embryonic brains were sectioned into 30 μm thick sections in the coronal plane using a Leica Cryostat CM1850 (Leica Microsystems). Sections were permeabilized with 0.3% Triton X100, then incubated with a blocking solution (3% normal goat serum prepared in PBS with 0.3% Triton X-100) and then incubated overnight with primary antibody prepared in blocking solution. An appropriate secondary antibody conjugated with an Alexa series fluorophore was used to detect the primary antibody. Draq5 (1:10,000 dilution, Cell Signaling) or 4′,6-diamidino-2-phenylindole (DAPI, 5 μg/ml, Invitrogen) were included in the secondary antibody solution to stain nuclei.

### IdU labeling and staining

Time-mated female mice were injected with 100 mg/kg IdU in PBS intraperitoneally at E14.5 or E15.5. Brain sections were prepared as described above in the Immunostaining section. Brain sections were permeabilized and DNA denatured with 4N HCl in PBS with 0.1% Triton X-100 for 30 minutes at 37°C, following by neutralizing with 0.1M sodium borate (pH 8.5) for 15 minutes at room temperature. Alternatively, sections were incubated with 0.5 unit/μl DNase I (TaKaRa, Kusatsu, Shiga, Japan) for 60 minutes at 37°C. After acid treatment or DNase I treatment, sections were subjected to the immunostaining procedure as described above in the Immunostaining section.

### Image acquisition and quantification

The dorsal view of post-fixed brain tissue was imaged by a FMA050 color CCD (AmScope, China). Epi-fluorescent images were taken by a DFC365FX monochrome CCD (Leica). Z-stack confocal images were acquired with a Leica SP8 confocal microscope. Specifically, Cux1^+^, PAX6^+^, and Tbr2^+^ cells were imaged with a 10X/NA0.75 or 20X/NA 0.7 objective, and the Z-stacks were taken at 0.5 μm intervals, 5 μm-total thickness were imaged and projected. BLBP^+^ processes were imaged with a 40X/NA 0.95 objective and the Z-stacks were taken at 0.5 μm intervals, 5 μm-total thickness was imaged and projected.

Triple-immunostaining to visualize NeuN^+^, S100β^+^, and IdU^+^ cells were simultaneously imaged with a 10X/NA 0.75 objective with a computational zoom of 0.75x and taken at 1μm intervals. Projection images of 5 μm-thickness were used to count cell numbers. The length, area, and numbers were quantified by using NIH ImageJ software. The whole cortex was divided into 10 bins (100 μm width per bin), and the cell counts were reported as the percentage of the total cell for each bin.

### Tissue clearing methodology

Brain samples were prepared as described before in the Immunostaining section. The fixed brains were sectioned into 1mm-thick sections in the coronal plane using a Leica VT-1000 vibrating microtome (Leica Microsystems). Brain sections were cleared using the Scale protocol ^50^, which preserves the red fluorescence signal expressed in this mouse line. Briefly, the brain sections were incubated in ScaleSQ(5) [22.5% d-sorbitol, 9.1M of urea, and 5% Triton X-100] for 2h in room temperature, followed by incubation in ScaleS4(D25) [40% d-sorbitol, 10% glycerol, 4M of Urea, 0.2% Triton X-100, 25% dimethyl sulfoxide] overnight at 4°C ^50^.

### Two-photon image acquisition and data analysis

The 1mm-thick cleared brain sections were mounted in 100 mm diameter Petri dishes filled with fresh ScaleS4(D25) solution and imaged for 15~20 hours. Images were acquired using a Nikon A1R MP+ multi-photon microscope (Nikon Instruments Inc., Melville, NY, USA) equipped with an InSight DeepSee infrared pulsed laser (Spectra-Physics Inc., Santa Clara, USA). The brain section was imaged with a 10X/NA 0.5 objective at 1μm intervals in the z-axis. Image analysis was performed utilizing the Surface module of Imaris v 9.2, (Bitplane Inc., Zurich, Switzerland). To quantify the volume of axonal projections in the striatal region, a 3D region of interest (ROI) measuring 800 μm × 800 μm × 700 μm (W×H×D) was selected from the most lateral portion of the striatum visible in the slice containing the anterior part of the hippocampus. The total volume occupied by the cortical glutamatergic axons (tdTomato^+^ axons) was extracted. To account for brain size differences in control and GOF brain, the total volume was further normalized to the area of the corresponding slice. The area of the slice was measured by manually contouring the middle plane of the z-stack image of interest. For the post commissural fornix analysis, a 3D ROI measuring 2850 μm × 1550 μm × 650 μm (W×H×D) was selected from the portion of the fornix immediately posterior to the anterior commissure. Measurements including volume, ellipticity prolate, and ellipticity oblate were extracted. The ellipticity prolate parameter represents the ROI’s similarity to a prolate ellipsoid. The ellipticity oblate parameter represents the ROI’s similarity to an oblate ellipsoid. The ellipticity values range from 0 to 1, being 1 a perfect ellipse (prolate or oblate) ^93^. To further investigate the characteristics of post commissural fornix, four horizontal 2-dimensional (2-D) planes were extracted from the acquired 3D volume containing the post commissural fornix, and the maximum diameter of fornix bundle was measured. To illustrate the axonal projections, we prepared a cartoon scheme (Fig. 7C, 9A) utilizing the Scalable Brain Atlas Composer ^94^.

### RNA-seq and pathway analysis

Cortices from E15.5 embryonic brains from control and GOF mice were used for RNA-seq experiments. Total RNA was extracted from brain tissue by RNeasy Mini Kit (Qiagen, Qiagen, Hilden, Germany) and followed by on-column DNase digestion according to the manufacturer’s instruction. RNA sequencing was performed by the Center for Medical Genomics, Indiana University School of Medicine. The concentration and quality of total RNA samples were first assessed using an Agilent 2100 Bioanalyzer. The RNA integrity number of samples was higher than 8.4 for all samples. Total RNA from different biological replicates (n=5 for control embryos, n=4 for GOF embryos) was used. Five hundred ng of RNA per sample was used to prepare a dual-indexed strand-specific cDNA library using TruSeq Stranded mRNA library Prep Kit (Illumina). The resulting libraries were assessed for their quality and size distribution using Qubind Agilent 2100 Bioanalyzer. One and a half picomoles of pooled libraries were sequenced in the 75bp single-end configuration on a NextSeeq500 (Illumina) using a NextSeq 500/550 High Output Kit. More than 90% of the sequencing reads reached Q30 (99.9% based call accuracy). The sequencing data was first assessed using FastQC (Babraham Bioinformatics, Cambridge, UK) for quality control. Then all sequenced libraries were mapped to the mouse genome (UCSC mm10) using STAR RNA-seq aligner. Genes with read counts per million (CPM) > 0.2 in more than 3 of the samples were kept. Differential expression analysis was performed using edgeR. False discovery rate (FDR) was computed from *p-values* using the Benjamini-Hochberg procedure. The multiple dimensional scaling (MDS) plot of RNA samples was drawn using plot MDS function in edgeR. The distance represented the leading log-fold-changes between each pair of RNA samples, which is the average (root-mean-square) of the largest absolute log-fold changes between each pair ^95^. This visualizes the differences between the expression profiles of different samples in two dimensions. Ingenuity Pathway Analysis (IPA, Qiagen, Germantown, MD) was performed for differentially expressed genes with FDR < 0.05. Enrichment of canonical pathways and disease and bio functions were identified with threshold p< 0.005. The activity of pathways and function was inferred as Z-scores. Positive z-score indicated increased activity, while negative z-score indicated inhibited activity.

## Supporting information

RNA-seq- Canonical Pathway analysis

RNA-seq DE list

## Data and statistical analysis

Data were analyzed using GraphPad Prism 7.04 software (GraphPad Software, San Diego, CA). In figures, data are expressed as means ± SEM. We employed the unpaired t-test and Two-Way ANOVA to examine data, as presented in figure legends.

## Acknowledgments

We thank Dr. Jean Hébert for *FGFR3*^*K650E*^ conditional mice and helpful comments from Drs. Chia-Shan Wu and Ken Mackie. We also thank Bruce Henry for technical assistance. Confocal images were taken in the Light Microscopy Imaging Center at Indiana University Bloomington. The sequencing was performed in the Center for Medical Genomics (CMG) at Indiana University School of Medicine, a sequencing core facility of the Indiana Clinical and Translational Sciences Institute.

## Author Contributions Statement

JY. H. and HC. L. designed research; JY. H., B. B. K, M. L. M., M. L. R., E. P. D., and J. M. G. performed research; JY. H., B. B. K, M. L. M., M. L. R., E. P. D., and J. M. G. analyzed data; JY. H., B. B. K, and HC. L. wrote the paper.

## Additional Information

The authors declare no competing financial interests.

This project was supported by the National Institutes of Health (NS048884 and NS086794 to H.C.L) and the Indiana Clinical and Translational Sciences Institute, funded in part by grant #UL1TR001108 from the National Institutes of Health, National Center for Advancing Translational Sciences, Clinical and Translational Sciences Award.

