## Supplementary Figures for "Enhanced FGFR3 activity in post-mitotic principal neurons during brain development results in cortical dysplasia and axon miswiring"

**Title**

### Supplementary Figure 1

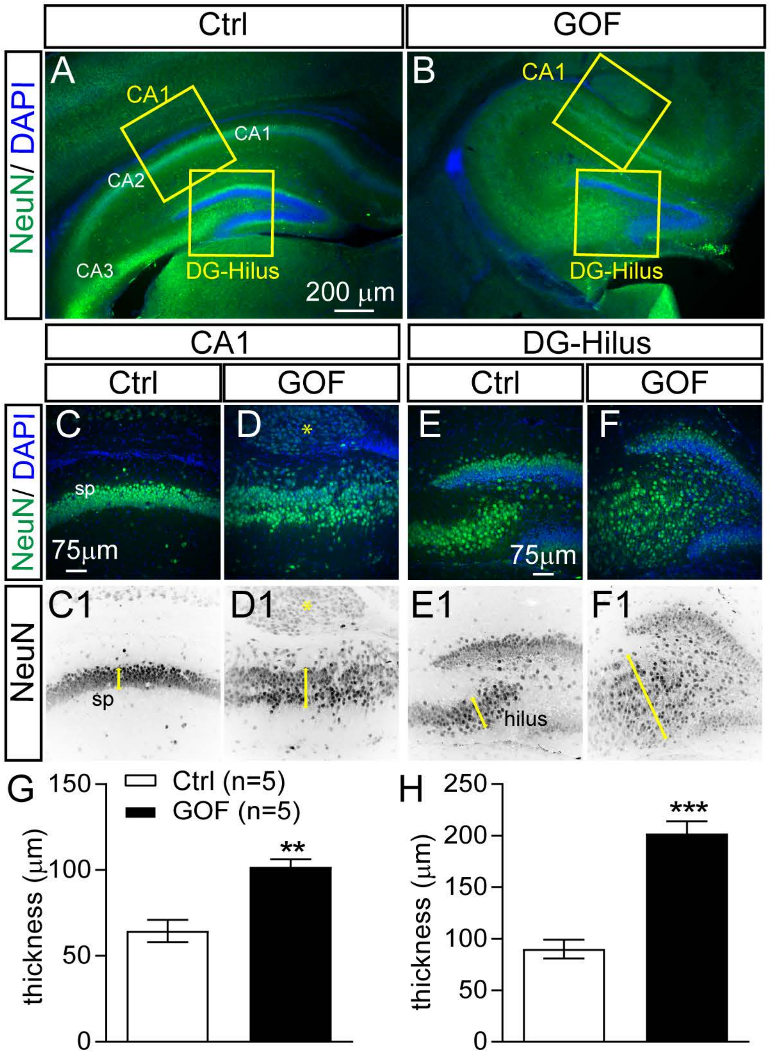

### Supplementary Fig 2

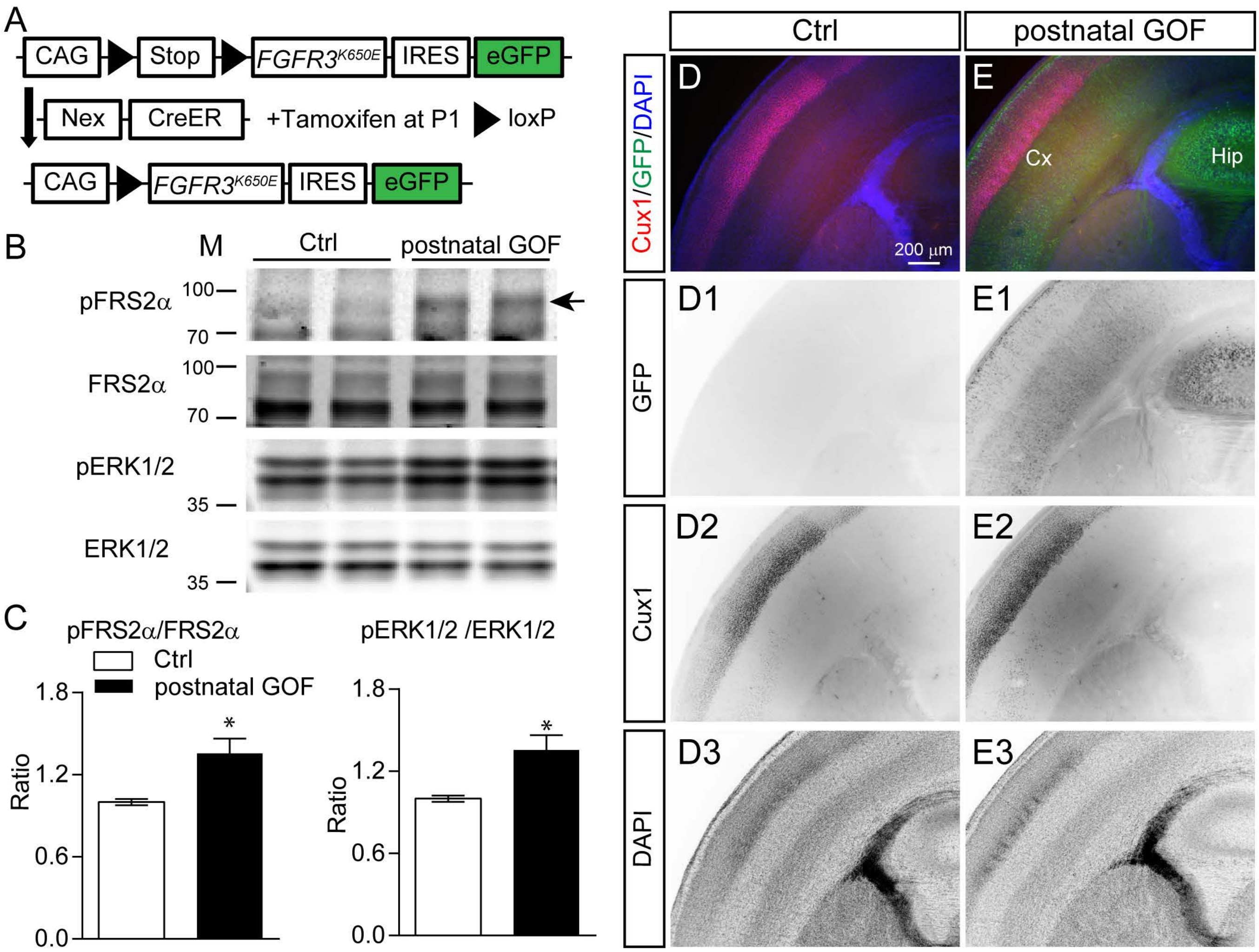

### Supplementary Figure 3

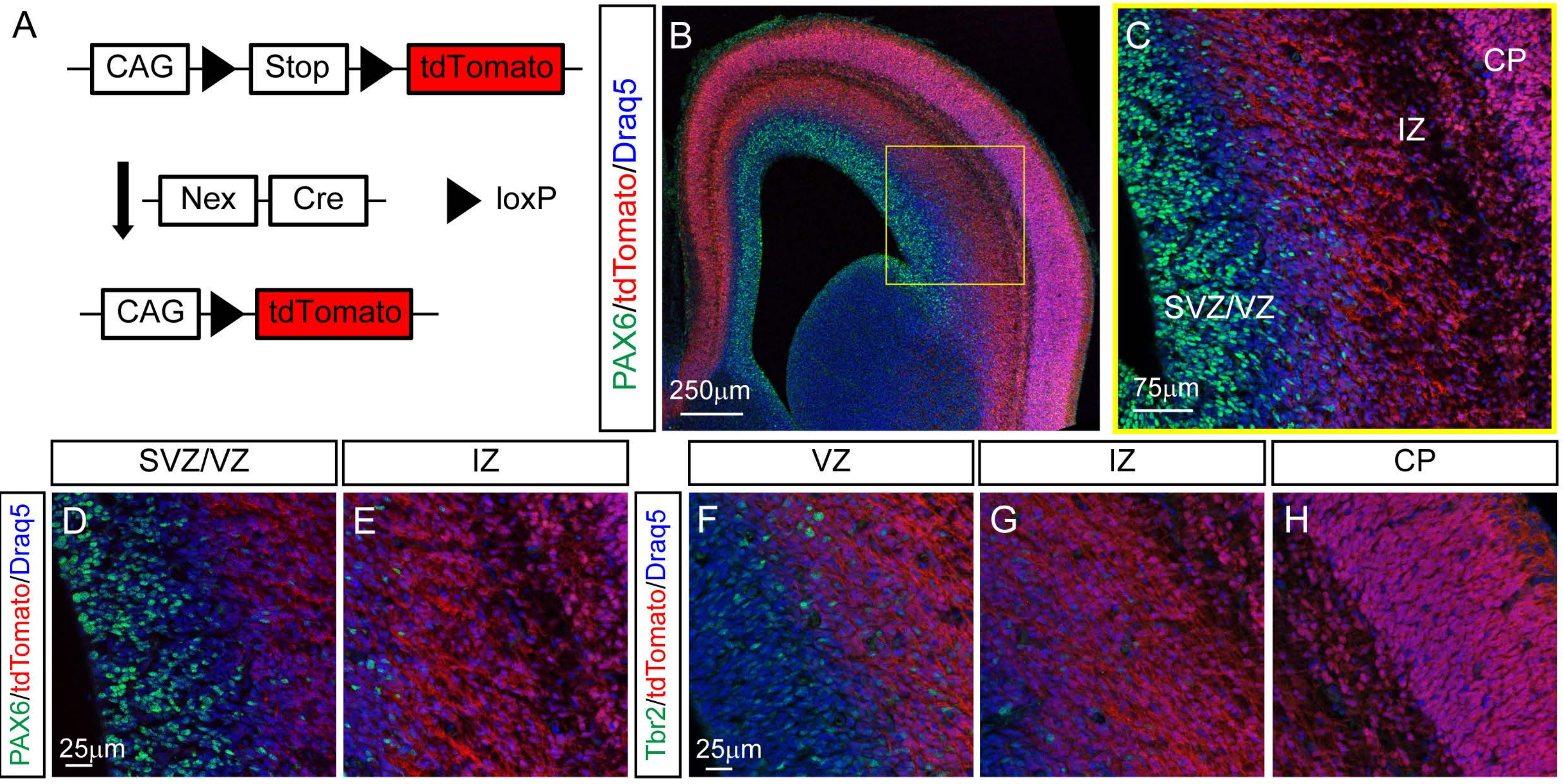

### Supplementary Figure 4

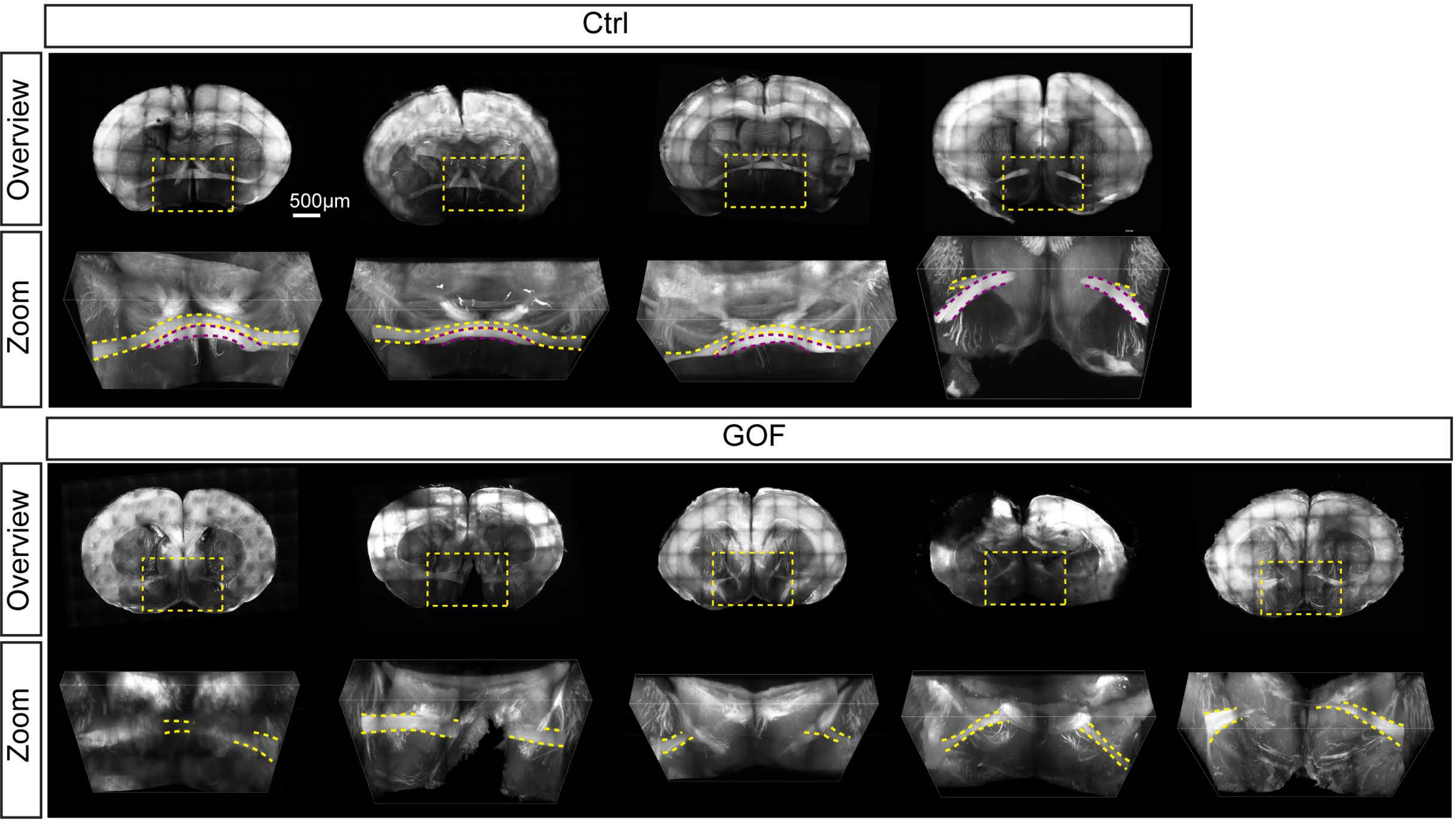

### Supplementary Figure 5

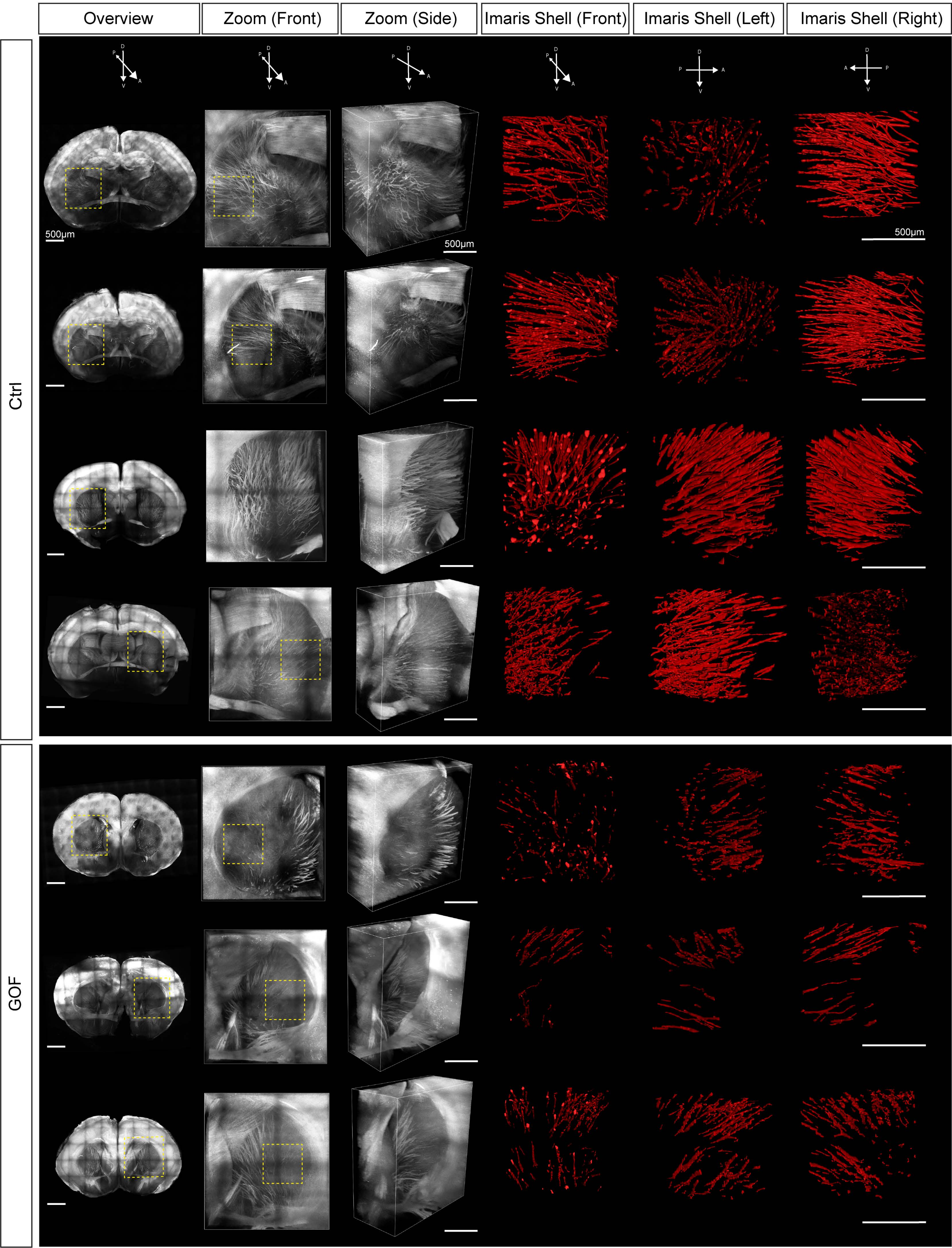

### Supplementary Figure 6

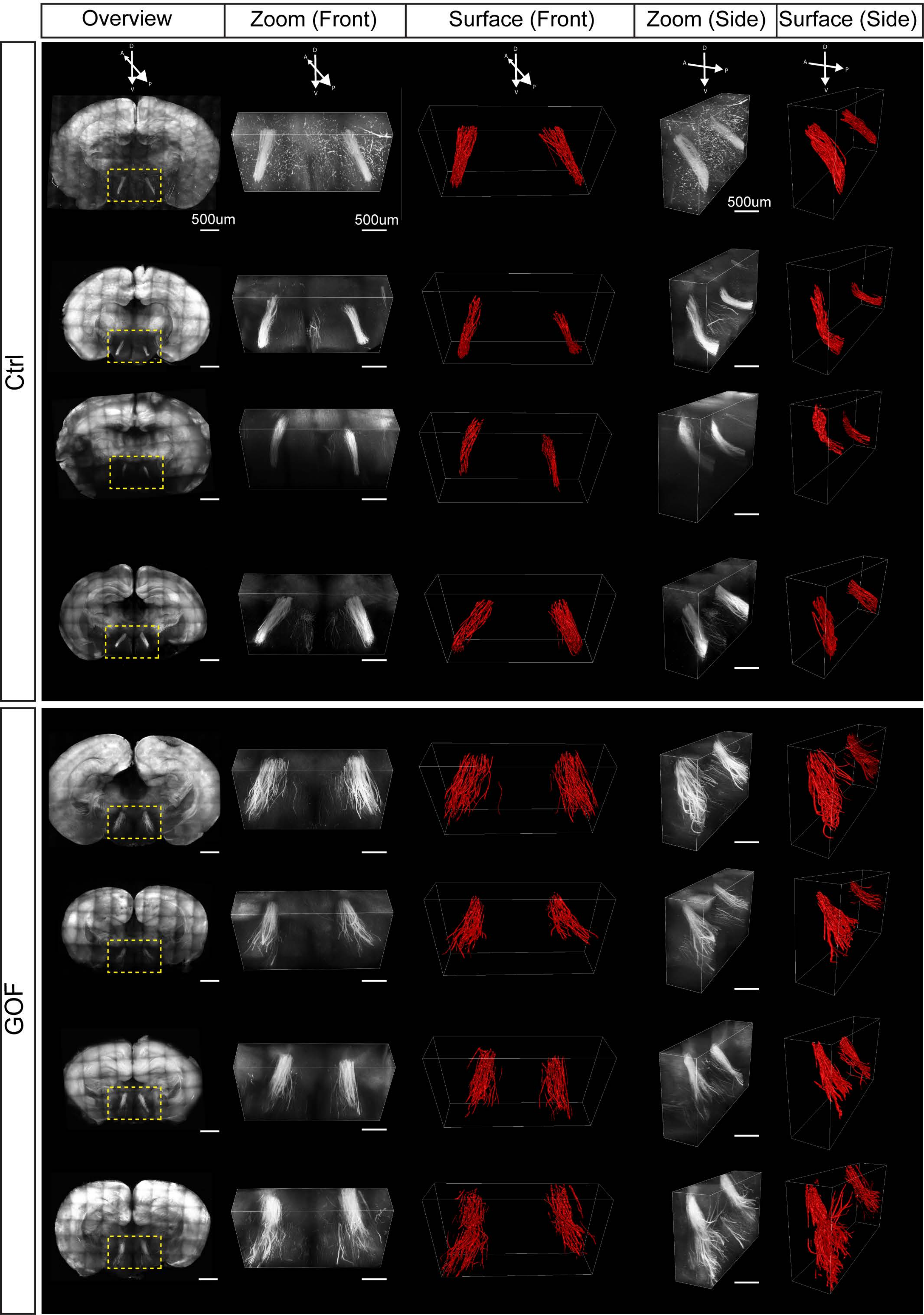

### Supplementary Figure 7

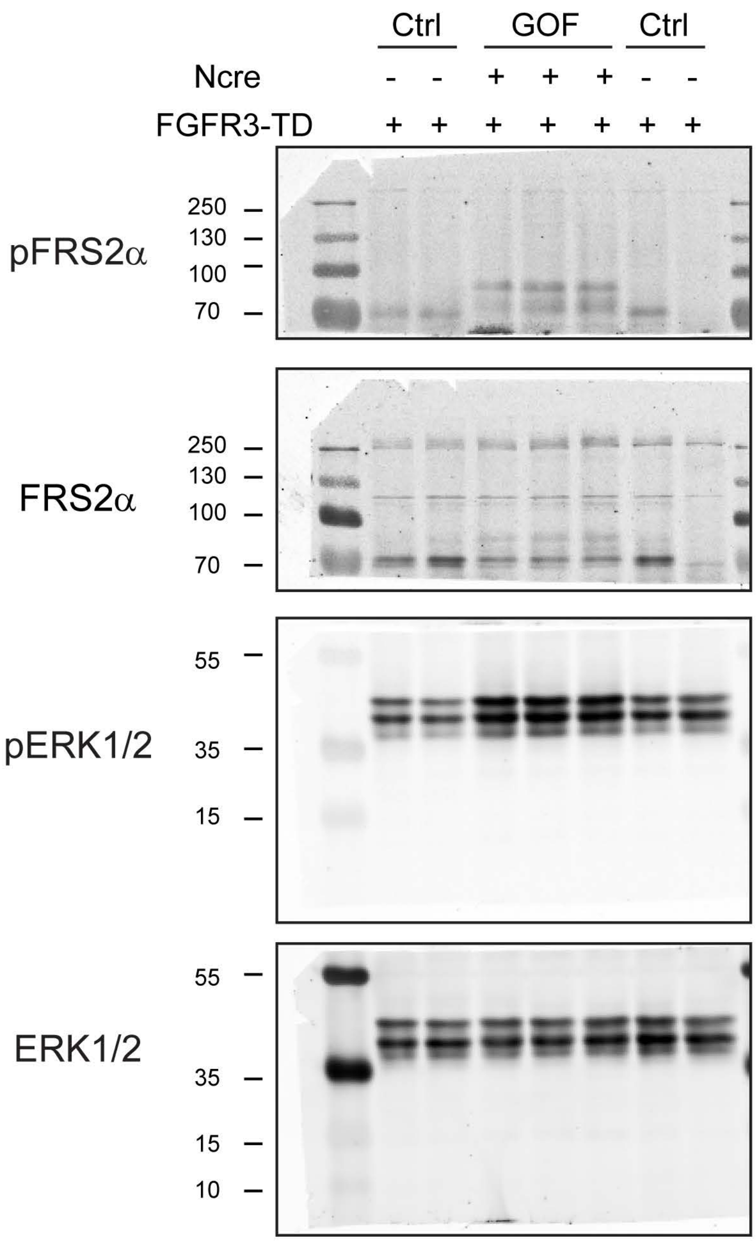

**Supplementary Figure 1. Expressing FGFR3$^{K650E}$ in NEX-lineage neurons results in aberrant hippocampal structure.** (A, B) NeuN staining of coronal sections showing the hippocampus in P7 control (Ctrl) and GOF brains. (C-F) Higher magnification images in CA1 and dentate gyrus (DG)-hilus (corresponding locations marked with yellow boxes in A and B). C1, D1, E1, and F1 show the inverted images of NeuN. Yellow star indicates a cluster of heterotopic neurons found in GOF brains. (G, H) Summary of the thickness of the NeuN$^{+}$ layer (indicated by the yellow line in C1-F1 (Ctrl, n=5; GOF, n=5) in CA1 (G) and DG-hilus (H). Student's-t test. **, $p < 0.01$; ***, $p < 0.001$.

**Supplementary Figure 2. Postnatal FGFR3 GOF did not perturb cortical laminations.** (A) The diagram shows how postnatal-GOF mice were generated. Tamoxifen (100 mg/Kg) was injected at P1 to activate CreER. Black triangles indicate loxP sites. (B) Western blots show the abundance of pFRS2α, FRS2α, pERK1/2, ERK1/2 in the S1 cortex of P7 ctrl and postnatal GOF mice. (C) Summaries for the fold changes of pFRS2α to FRS2α and pERK1/2 to ERK1/2 (ctrl, n=5; GOF, n=5) in GOF mice. Student's-t test. * $p<0.05$. (D-E) GFP and Cux1-staining in the P7 S1 cortex of ctrl and postnatal-GOF brains. D1-D3 and E1-D3 show the inverted images of the indicated channel. Cx, cortex; Hip, hippocampus.

**Supplementary Figure 3. tdTomato reporter mice demonstrate that majority of Cre mediated recombination occurs in post-mitotic neurons.** (A) The diagram shows how Nex-cre;TdTomato mice were generated. (B, C) Images show tdTomato and Pax6 double staining in E15.5 brains. C is the high magnification view of yellow box in B. (D, E) Images for PAX6 and tdTomato double staining in sub-ventricular zone (SVZ)/ventricular zone (VZ) and intermediate zone (IZ). (F-H) Tbr2 and tdTomato staining in VZ, IZ, and cortical plate (CP).

**Supplementary Figure 4. GOF mice have no olfactory limb of the anterior commissure**. More images from different animals to show that the olfactory limb of the anterior commissure is missing in GOF mice.

**Supplementary Figure 5. FGFR3 GOF mice have reduced numbers of axonal projections in the striatum.** More images from different animals. D, dorsal; V, ventral, A, anterior; P, posterior.

**Supplementary Figure 6. FGFR3 GOF disrupts axonal fasciculation of postcommissural fornix.** More images from different animals. D, dorsal; V, ventral, A, anterior; P, posterior.

**Supplementary Figure 7.** Full-length blots of Figure1.
